# Phase separation of Myc differentially modulates the transcriptome

**DOI:** 10.1101/2022.06.28.498043

**Authors:** Junjiao Yang, Chan-I Chung, Jessica Koach, Hongjiang Liu, Ambuja Navalkar, Qian Zhao, Xiaoyu Yang, Liang He, Tanja Mittag, Yin Shen, William A. Weiss, Xiaokun Shu

**Affiliations:** Department of Pharmaceutical Chemistry, University of California, San Francisco, San Francisco, California, USA; Cardiovascular Research Institute, University of California, 1,2California, USA; Departments of Neurology, Neurological Surgery, Pediatrics, and Helen Diller Family Comprehensive Cancer Center, University of California, San Francisco, San Francisco, CA, USA; Institute for Human Genetics, Departments of Neurology, Weill Institute for Neurosciences, University of California, San Francisco, San Francisco, CA, USA; Department of Structural Biology, St. Jude Children’s Research Hospital, 262 Danny Thomas Place, Memphis, TN 38105, USA

**Author notes:** These authors contributed equally (Junjiao Yang, Chan-I Chung). Correspondence to: Xiaokun Shu.

## Abstract

Dysregulation and enhanced expression of *MYC* transcription factors (TFs) including *MYC* and *MYCN* contribute to the majority of human cancers. For example, *MYCN* is amplified up to several hundred-fold in high-risk neuroblastoma. The resulting overexpression of N-myc aberrantly activates genes that are not activated at low N-myc levels and drives proliferation and cell survival. Whether increasing N-myc levels simply mediate binding to lower-affinity binding sites in the genome or fundamentally changes the activation process remains unclear. One such activation mechanism that could become important above threshold levels of N-myc is the formation of aberrant transcriptional condensates through phase separation. Phase separation has recently been linked to transcriptional regulation, but how strongly it contributes to gene activation remains unclear. Here we characterized the phase behavior of N-myc and showed that it can form dynamic condensates that bear the hallmarks of transcriptional activity. We tested the contribution of phase separation to N-myc-mediated gene expression by using a chemogenetic tool that allowed us to compare non-phase-separated and phase-separated conditions at identical N-myc levels, which both showed a strong impact on gene expression compared to no N-myc expression. However, we found that only a small fraction of <3% of N-myc-regulated genes is further affected by phase separation, but that these events include the activation of key oncogenes and the repression of tumor suppressors. Indeed, phase separation increases cell survival by ∼15% corroborating the biological effects of the transcriptional changes. However, our results also show that >97% of N-myc-regulated genes are not affected by N-myc phase separation, highlighting that transcription can be activated effectively by diffuse complexes of TFs with the transcriptional machinery.

## Introduction

*MYC* family transcription factors are major contributors to human tumorigenesis. Expression of Myc is deregulated and enhanced in many types of cancers due to copy number changes, chromosomal translocations, or upstream oncogenic signaling ^3–6^. For instance, *MYCN* is highly amplified 100-to-300 fold in nearly half of high-risk neuroblastoma patients ^7–10^. While upregulated Myc expression induces tumor development in many tissues, depletion of Myc abolishes tumorigenesis and results in tumor regression in various tumor models ^11–15^. Given that high Myc expression does not only enhance the transcriptional activation but changes the pattern of gene expression to promote proliferation and cell survival, one important question is how the changes to the transcriptional program are caused by rising Myc levels. Do increasing Myc concentrations allow binding to progressively lower-affinity binding sites in the genome ^16^, or does the activation mechanism fundamentally change?

One potential mechanism for a fundamental change in activation mechanism is the formation of transcriptional condensates through phase separation (PS). Phase separation mediates the formation of a dense phase above a threshold concentration of biomolecules ^17–21^. Recently, many transcription factors (TFs) have been reported to undergo PS and form biomolecular condensates (also known as membraneless compartments, granules, or liquid droplets) when protein concentration surpasses a threshold concentration ^22–25^. Biomolecular condensates compartmentalize interacting proteins and signaling complexes in percolated dense phases ^18–20,25–28^. Condensates of transcription factors have been proposed and demonstrated to compartmentalize transcriptional machinery ^22–25,29^. However, whether phase separation really changes transcriptional output remains a matter of discussion ^30^. Answers to this question are hampered by conceptual and technical challenges. Mutations that change phase behavior are employed in many studies to correlate the driving force for phase separation with function ^23,31^, but these mutations likely also impact the ability of TFs to form diffuse complexes with the transcriptional machinery ^22,32^. Such diffuse complexes form via transient multivalent interactions as characterized in detail for the yeast transcription factor GCN4 ^33^. These same interactions are also conducive to mediating the formation of the three-dimensional network structure spanning condensates. Furthermore, associative biomolecules that drive phase separation are expected to form higher-order complexes, or so-called pre-percolation clusters, below the saturation concentration ^34^. It thus remains an open question whether the transcriptional output observed in the presence of transcriptional condensates would also be achieved without phase separation. Interestingly, a recent study shows that phase separation does not enhance transcription^35^, though this study changed TF expression levels to compare activity in cells with and without condensates near the threshold concentration of PS. But assigning the threshold level for the formation of condensates on chromatin is challenging under cellular context where heterotypic multicomponent interactions result in non-fixed saturation concentration ^36^. Furthermore, changing TF levels will itself influence activity. Given that aberrant transcriptional condensates may offer attractive opportunities to develop new therapeutics ^37,38^, these challenges in understanding the role of phase separation in transcription call for new tools that allow us to assess transcriptional activity upon turning on phase separation without changing expression levels or making mutations.

Here, we develop a chemogenetic tool which we use to turn on N-myc phase separation in a neuroblastoma cell line and compare gene expression to that of cells with equi-concentrated N-myc without phase separation. We find that N-myc condensates are transcriptionally active and largely regulate gene expression similar to diffuse N-myc. However, a small fraction of genes is altered further upon phase separation, and these may be important for Myc-dependent oncogenesis. Our work thus shows a role for phase separation of overexpressed transcription factors in disease but simultaneously shows that phase-separated TFs do not necessarily mediate all of the enhanced functions.

## RESULTS

### N-myc undergoes phase separation in cells

To test whether N-myc forms condensates, we used immunofluorescence imaging in *MYCN*-amplified human Kelly neuroblastoma cells. This indicated that N-myc protein formed puncta in the nucleus (Fig. 1a), which were not observed upon treatment with a MYC/MAX dimerization inhibitor that was previously shown to degrade Myc ^39^ (Supporting Fig. S1). Importantly, in *MYCN*-nonamplified SH-EP neuroblastoma cells, we did not observe obvious punctate structures (Fig. 1a), consistent with the western blot analysis showing no N-myc expression in SH-EP (Supporting Fig. S2). To characterize the phase behavior of N-myc, we expressed the fusion protein N-myc-mEGFP in SH-EP cells that did not express N-myc and conducted live-cell imaging. N-myc-mEGFP was evenly distributed and did not form punctate structures until its expression level was above a threshold concentration (i.e. the saturation concentration; Fig. 1b).

**Fig. 1.**
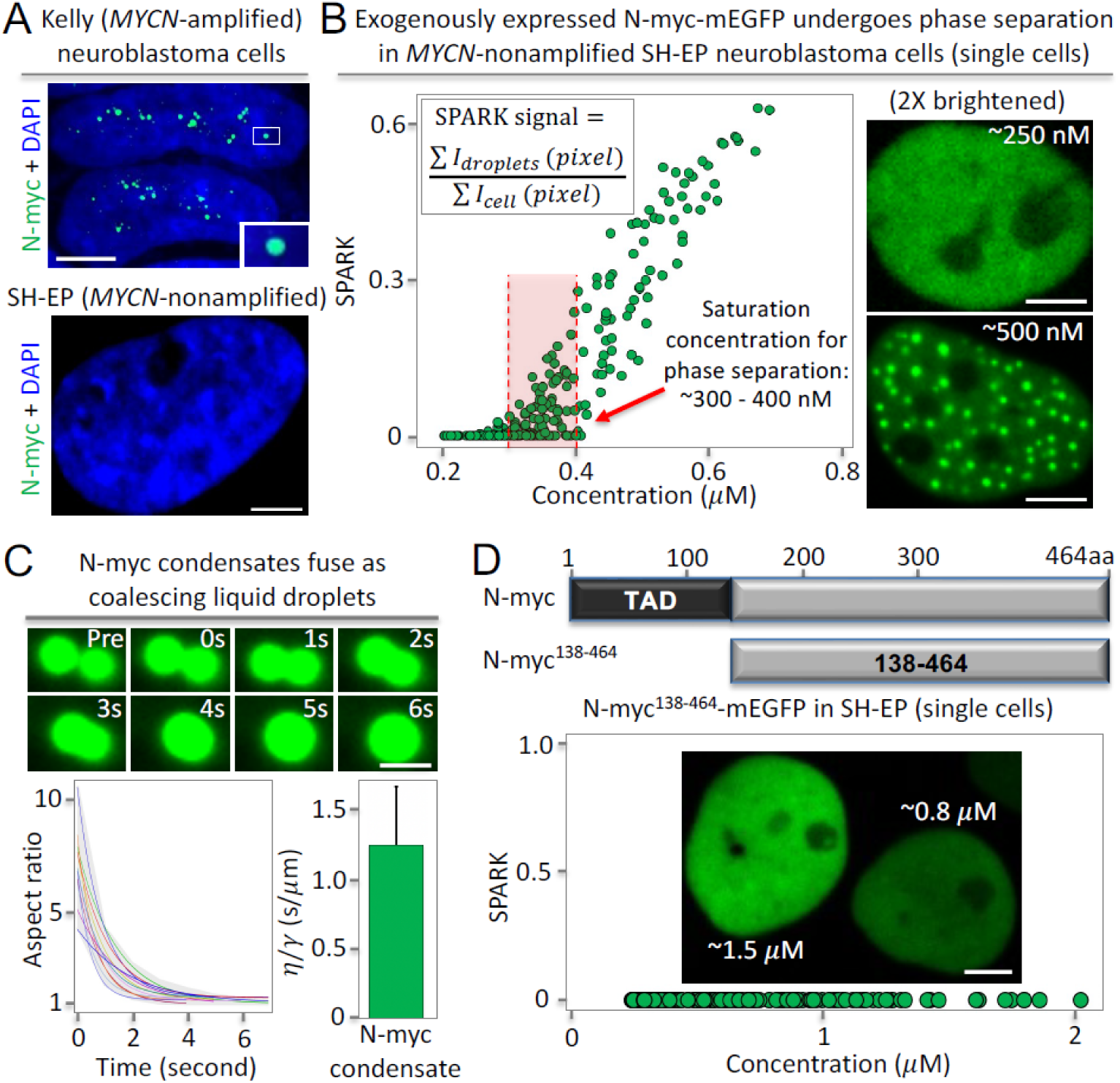
N-myc undergoes phase separation, which requires the intrinsically disordered transactivation domain. (**A**) Immunofluorescence images of N-myc in *MYCN*-amplified neuroblastoma Kelly cells (top panel) and the *MYCN*-nonamplified SH-EP cells (bottom panel). N-myc forms puncta in the Kelly cells but not in the SH-EP cells that express no or little N-myc (Fig. S3). Scale bar, 10 μm. (**B**) Expression of mEGFP fused N-myc in the neuroblastoma SH-EP cells that have no or little endogenous N-myc protein expression. Left: quantitative analysis of N-myc puncta formation against its protein level in single cells. Each green circle corresponds to an individual cell, i.e. ∼300 cells were analyzed. The concentration of the fusion protein was estimated based on purified mEGFP (see details in Methods). The red dashed lines depict a range of saturation concentrations for phase separation. Right: representative fluorescence images; the upper panel is brightened, the lower is not. Scale bar, 5 μm. (**C**) Fusion events between N-myc condensates. Top: fluorescence images of time course. Scale bar, 1 μm. Bottom-left: quantitative analysis of the fusion events. Individual lines represent the best fit line for individual fusion events. Gray shade shows the range of aspect ratio values from all events. Bottom-right: mean inverse capillary velocity extracted from fusion events (n = 12). Error bar represents standard deviation. (**D**) Quantitative analysis of the truncated N-myc lacking N-terminal TAD, which is an IDR. Each green circle corresponds to an individual cell (∼200 cells). Scale bar, 5 μm.

Our quantitative analysis yielded the percentage of N-myc in punctate structures. The percentage is dependent on protein expression levels, and it was ∼30 – 50% at N-myc level of 500 nM. We estimated the protein concentration in single cells based on purified mEGFP and were able to determine that the range (or confidence interval) of saturation concentrations of N-myc was ∼ 300 – 400 nM (see Methods and Supporting Fig. S3). Our data shows an interval rather than a single threshold concentration, consistent with previous report that under cellular context, heterotypic phase separation results in non-fixed saturation concentrations ^36^. Below the range of saturation concentrations, e.g. at ∼ 250 nM, N-myc was evenly distributed in the nucleus (Fig. 1b, upper-right). Above the saturation concentrations, e.g. at ∼500 nM, N-myc formed puncta in the nucleus (Fig. 1b, lower-right). Thus, our data show that N-myc-mEGFP undergoes a concentration-dependent spatial rearrangement. We estimated that the N-myc concentration is around 0.7 – 1 μM in the Kelly cells (Supporting Fig. S4). We characterized the relationship of number and size of the N-myc puncta to the protein levels and observed that the number of N-myc puncta increases as N-myc protein level increases (Supporting Fig. S5A). The size of N-myc puncta is in the range of 0.4 – 1 μm (diameter), and this distribution is independent of the protein levels (Supporting Fig. S5B). This suggests that N-myc tends to form new puncta when the protein level increases.

Next, we determined whether the N-myc puncta exhibit liquid-like properties. We conducted time-lapse imaging and observed that puncta can fuse and coalesce within a few seconds. The fusing puncta initially formed a dumbbell shape, which over time relaxed to a spherical shape (Fig. 1c). Quantitative analysis showed that the aspect ratio of the fusing puncta over time fits well to a single exponential curve (Fig. 1c, lower left), which is a well-known characteristic of coalescing liquid droplets ^40,41^. We used the data to determine the inverse capillary velocity (= η/γ; here γ is surface tension of the droplet; η is viscosity), which was 1.25 ± 0.42 (s/μm) (Fig. 1c, lower right). Thus, the N-myc puncta are liquid-like condensates formed by phase separation when the concentration of N-myc-mEGFP exceeds threshold levels, which are reached by many *MYCN*-amplified cancer cell lines.

N-myc, which encompasses 464 residues, has an intrinsically disordered transactivation domain (TAD), which encompasses conserved N-terminal motifs including three “Myc boxes” (MB0-II) from residues 1 - 137 residues ^42,43^. To examine the role of the TAD in N-myc PS, we designed and characterized a TAD truncation mutant (N-myc^138–464^). Live cell imaging revealed that this mEGFP-tagged fusion protein (N-myc^138–464^–mEGFP) no longer formed condensates even above 2 μM concentration (Fig. 1d), ∼ 5-fold above the threshold concentration of PS for full-length N-myc. Therefore, our data demonstrate that N-myc PS requires the intrinsically disordered TAD, consistent with PS of many other proteins that also rely on their IDR ^19^.

### N-myc condensates contain DNA-binding partner MAX and localize to target gene loci

To examine whether the N-myc condensates are transcriptionally active, we first asked whether N-myc condensates contain the obligatory DNA-binding and dimerization partner MAX. To visualize MAX in living cells, we labeled it with the red fluorescent protein mKO3. Multicolor fluorescence imaging showed that MAX also had a punctate appearance, and that MAX and N-myc puncta overlapped (Fig. 2A). In cells without N-myc-mEGFP, MAX did not have a punctate appearance (Supporting Fig. S6). These data suggest that N-myc condensates recruit its DNA-binding partner MAX or that N-myc and MAX co-condense.

**Fig. 2.**
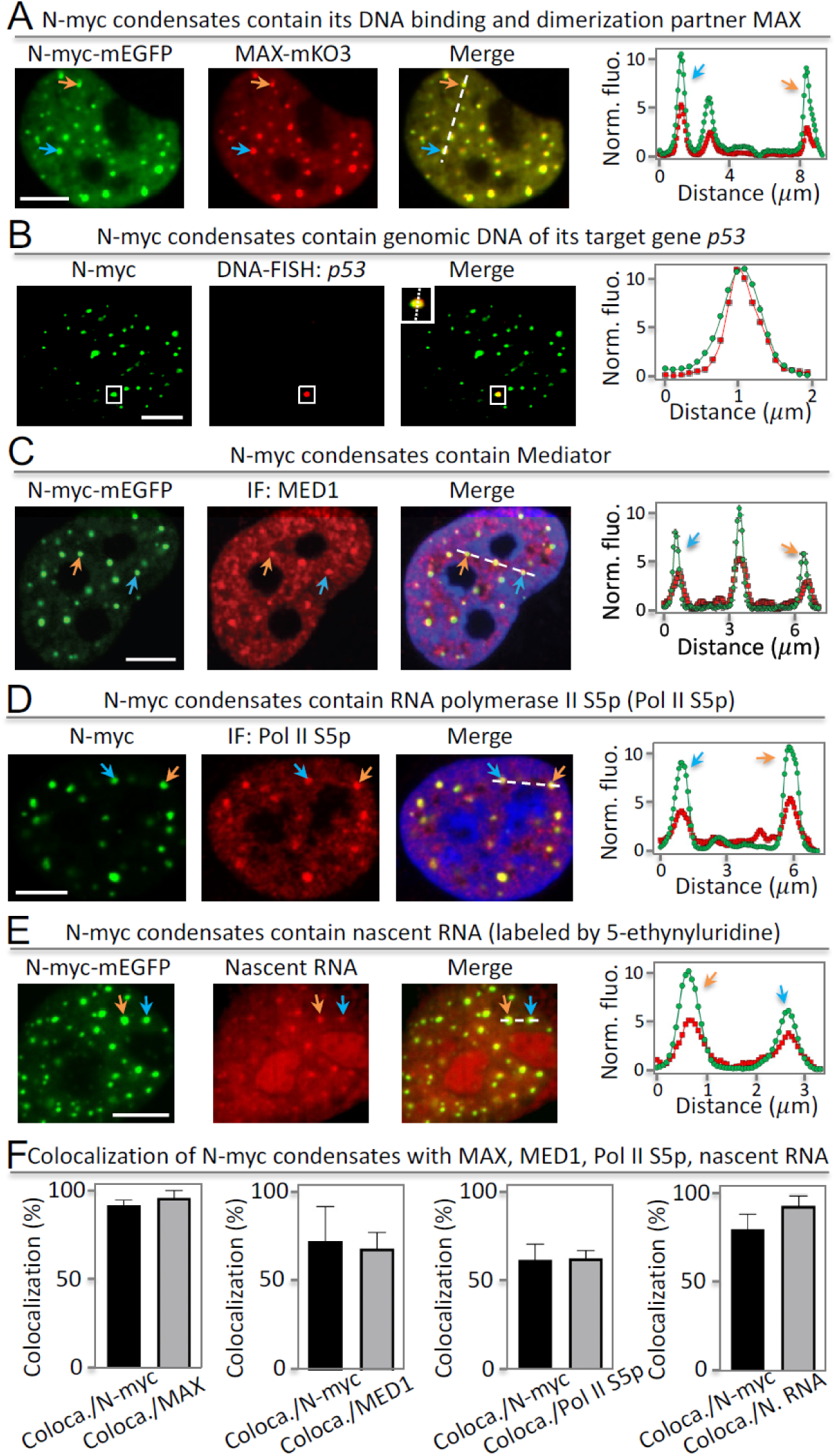
N-myc condensates contain DNA-binding and dimerization partner, genomic DNA, transcriptional machinery and nascent RNA. (**A**) Fluorescence images of N-myc-mEGFP and MAX-mKO3 in SH-EP cells. The arrows point to representative condensates. The fluorescence intensity profile (right panel) is extracted from the position shown by the dashed line. Colocalization in example condensates is indicated by arrows. (**B**) Fluorescence images of N-myc condensates with single molecule DNA FISH against *p53*. (**C**) Fluorescence images of N-myc condensates with immunofluorescence (IF)-imaged MED1. (**D**) Fluorescence images of N-myc condensates with IF-imaged Pol II S5p. (**E**) Fluorescence images of N-myc condensates with nascent RNA labeled by 5-ethynyluridine. (**F**) Percentage of N-myc condensates that colocalize with puncta of other components. The percentage is determined by the ratio of coloca./N-myc = number of colocalized condensates between N-myc and MAX divided by number of N-myc condensates. Equivalent analysis for other pairs is also shown. Data are mean ± SD (n = 13 cells). Scale bars, 5 μm (A – E).

To test whether N-myc condensates localized to DNA loci of an example N-myc target gene, we chose *p53* ^44,45^ and labeled the *p53* DNA locus using fluorescence in situ hybridization (FISH). Confocal fluorescence imaging revealed that N-myc condensates were associated with the genomic DNA locus of *p53* (Fig. 2B).

### N-myc condensates contain transcriptional machinery and nascent RNA

To further test whether N-myc condensates may be transcriptionally active, we determined whether they contained transcriptional machinery, including the Mediator and RNA polymerase II (Pol II). First, immunofluorescence imaging showed that the Mediator of RNA polymerase II transcription subunit 1 (MED1) had punctate localization in cells, consistent with previous studies ^46^. Furthermore, N-myc-positive condensates were a subset of MED1 condensates (Fig. 2C), indicating that N-myc condensates contain MED1. Second, we stained the cells with antibodies against phosphorylated Pol II at Ser5 (Pol II S5p) at the C-terminal domain. Immunofluorescence imaging showed punctate structures of Pol II S5p, which colocalized with N-myc condensates based on two-color imaging (Fig. 2D). Therefore, our data indicate that N-myc condensates contain the transcriptional machinery.

We next tested whether the N-myc condensates contained nascent RNA. We incubated cells with the uridine analog 5-ethynyluridine (EU) for 1 hour so that EU was incorporated into newly transcribed RNA. The EU-labeled nascent RNA was detected through a copper (I)-catalyzed cycloaddition reaction (i.e. “click” chemistry) using azides labeled with red fluorescent dyes ^47^. Fluorescence imaging revealed several punctate structures (Fig. 2E), many of which colocalized with the N-myc condensates, suggesting that these N-myc condensates contain nascent RNAs.

Lastly, we quantified the colocalization of N-myc condensates with MAX, MED1, Pol II S5p and nascent RNAs (Fig. 2F, Methods). We calculated that ∼92% of N-myc condensates contained MAX. The percentage of N-myc condensates that contain MED1, Pol II S5p and nascent RNAs is ∼ 72%, 62%, 80%, respectively. In aggregate, our data show that N-myc condensates have the hallmarks of transcriptionally active assemblies.

### N-myc condensates are dynamically regulated during cell mitosis

Biomolecular condensates have emergent properties that small complexes do not have, and these include ripening and coalescence. Coalescence of chromatin-bound condensates have the potential to bring distant genomic loci into proximity and potentially influence transcriptional programs ^32^. Hence the timescales on which they assemble and disassemble are expected to influence their function. Previous studies show that many biomolecular condensates disassemble during mitosis ^48^. Here we examined whether N-myc condensates were also regulated dynamically during the cell cycle. Live-cell fluorescence imaging showed that N-myc condensates dissolved when cells entered mitosis (Fig. 3A, left panel). Upon mitotic entry, chromatin condenses even though nuclear chromatin is already compacted in the interphase. It has been well established that many transcription factors disengage from chromatin when cells enter mitosis^49,50^. We thus decided to investigate the relationship between N-myc condensate dissolution and chromatin condensation upon entry into mitosis. To visualize chromatin, we labeled histone 2B (H2B) with a near-infrared fluorescent protein mIFP ^51^. This allowed us to quantify the volume of chromatin using fluorescent protein-labeled H2B ^52^. Time-lapse imaging revealed that the dissolution of N-myc condensates preceded chromatin condensation by ∼ 6 minutes (Fig. 3A, right panel). The dissolution of N-myc condensates also occurred before nuclear envelope breakdown (Fig. 3A, T ∼ 18 min.).

**Fig. 3.**
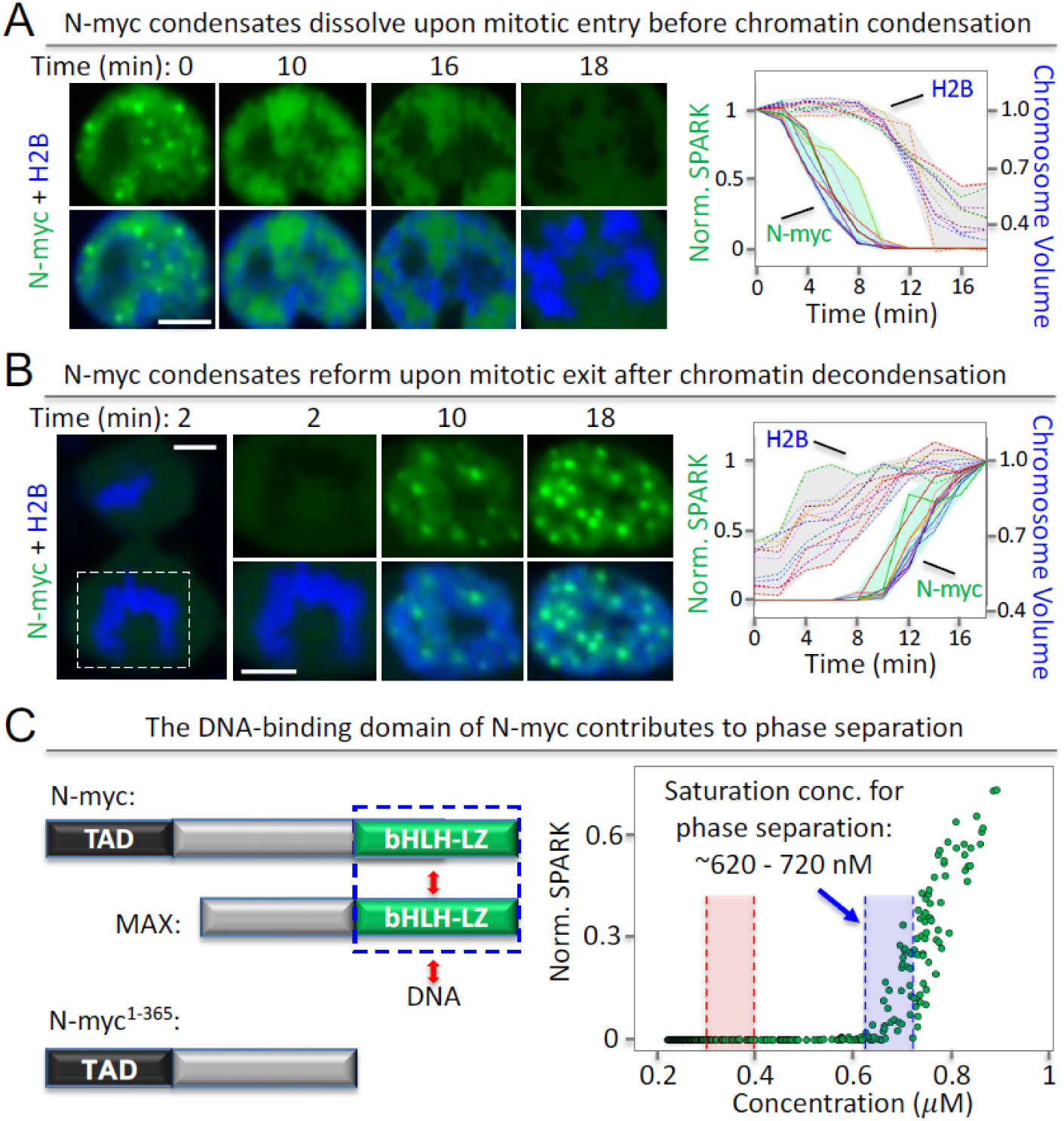
N-myc condensates dynamically disassemble and re-assemble upon mitotic entry and exit. (**A**) Time-lapse images of SH-EP cells expressing N-myc-mEGFP upon mitotic entry. The cells co-expressed mIFP-tagged histone 2B (H2B, in blue). Chromosome volume was calculated based on mIFP-H2B fluorescence. Right panel: quantitative analysis of N-myc condensate dissolution and chromosome condensation as a function of time. Each line represents single cell traces (n = 11 cells). (**B**) Time-lapse images of SH-EP cells expressing N-myc-mEGFP upon mitotic exit. Right panel: quantitative analysis of N-myc condensate reformation and chromosome de-condensation. Each line represents single cell traces (n = 12 cells). (**C**) Phase diagram of the truncated N-myc lacking the DNA-binding domain. The blue box depicts a range of saturation concentrations for N-myc^1–365^ phase separation. The red box depicts a range of saturation concentrations for full length N-myc phase separation (see Fig. 1B). Scale bars, 5 μm (A, B).

Next, we examined whether N-myc reformed condensates when cells exit mitosis. Time-lapse imaging revealed that indeed upon mitotic exit, N-myc condensates reappeared. We also observed that chromatin decondensed during mitotic exit, consistent with previous studies ^52^. Interestingly, during mitotic exit, N-myc condensates formed after chromatin decondensation with a delay of ∼ 6 minutes (Fig. 3B). This mirrors mitotic entry, where the dissolution of N-myc condensates occurred before chromatin condensation.

Our study thus reveals that N-myc condensates are dynamically regulated during mitosis, and that condensate disassembly and reassembly is correlated with chromatin condensation and de-condensation, respectively. Because many transcription factors disengage from chromatin when cells enter mitosis and re-associate with chromatin when cells exit mitosis ^49,50^, we investigated a potential role of the N-myc DNA-binding domain bHLH-LZ (366-464 aa) on N-myc phase separation. We truncated bHLH-LZ and measured phase separation of this truncation mutant N-myc^1–365^. Indeed, the range of saturation concentrations of this mutant is ∼ 620 – 720 nM (Fig. 3C, blue box), which is ∼ 2-fold more than that of full-length N-myc (∼ 300 – 400 nM, Fig. 1B; red box in Fig. 3C). Our data thus suggest that the DNA-binding domain contributes to N-myc phase separation.

### E-box DNA enhances condensate formation of purified N-myc

Given that we found a contribution of the DNA-binding domain to N-myc condensate formation, we sought to understand quantitatively how phase separation of N-myc is modified by binding to different DNA sequences. We thus turned to experiments in vitro with purified proteins and defined DNA oligonucleotides. CD spectroscopy showed that recombinant N-myc had a partially helical secondary structure as expected ^53^ (Fig. 4A). We refolded N-myc and Max independently and mixed them at a molar ratio of 10:3 (to disfavor Max/Max dimer formation), and the resulting CD spectra were comparable to the sum of the CD spectra of the individual components. N-myc/MAX bound to a DNA oligomer containing a single E-box (CACGTG in 26-mer, called 1Ebox DNA from hereon) with a *K*_d_ of ∼150 nM, indicating that our samples contained high-quality N-myc/Max dimers (Fig. 4B).

**Fig. 4.**
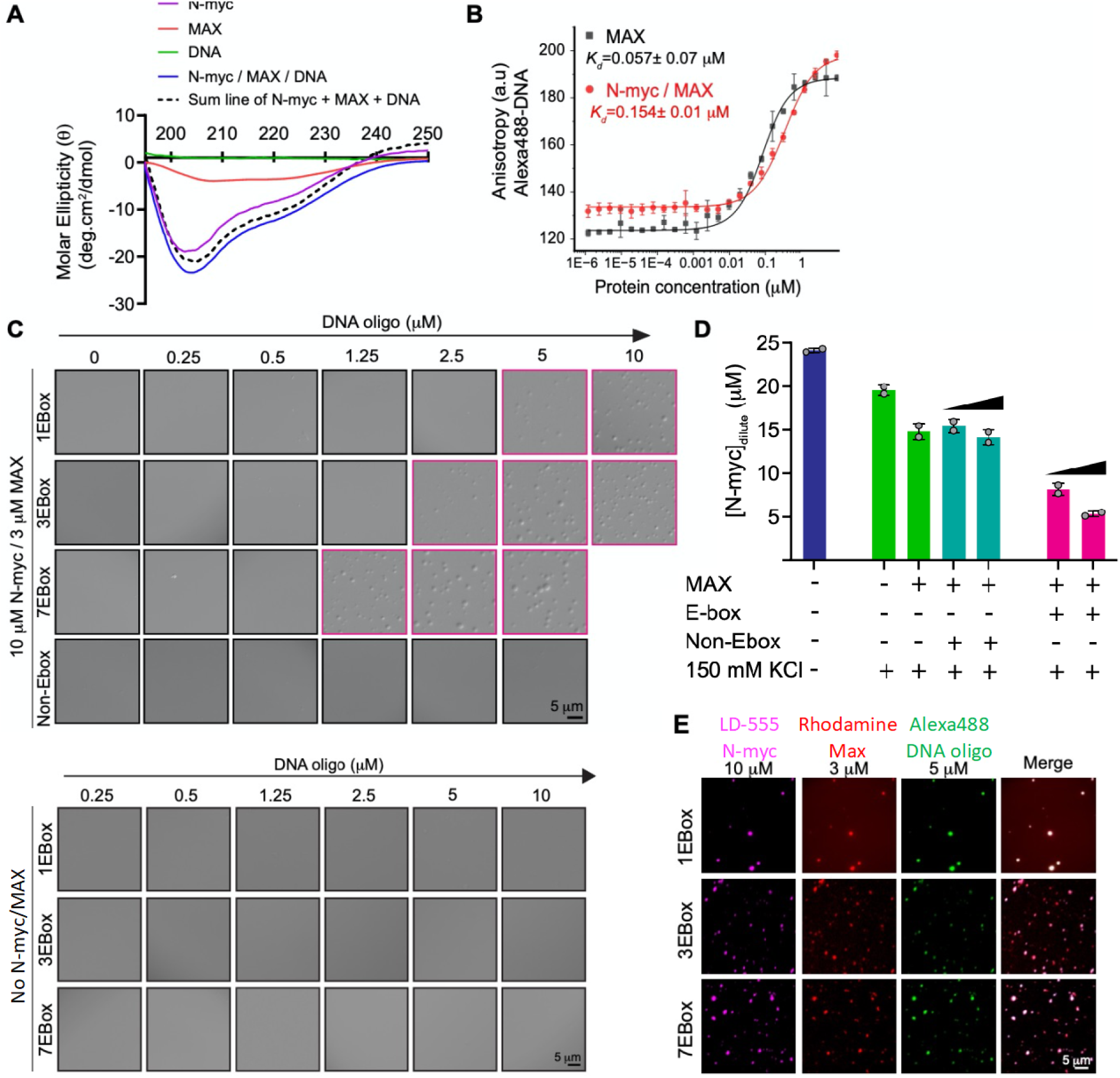
Phase separation of full-length N-myc protein is enhanced by Max and DNA oligonucleotides containing canonical Myc E-box sequences. **(A)** Far UV circular dichroism (CD) spectra for N-myc, MAX, oligonucleotide DNA containing 1 E-box sequence (CACGTG, referred to as 1EBox DNA) and for a complex of all three. The sum of the individual spectra is shown for comparison. The minima at 208 and 222 nm indicate helical structure. **(B)** Fluorescence anisotropy assay showing binding of the N-Myc / MAX heterodimer to 50 nM fluorophore-labeled 1Ebox DNA. The ratio of Myc / Max was fixed at 3:1. The data are the mean of 3 replicates, the error bars ± SD. **(C)** Upper panel: Differential Interference Contrast (DIC) microscopy images of N-myc / Max complex (10 µM Myc / 3 µM Max) in presence of 1Ebox (26-mer), 3Ebox (78-mer) and 7Ebox (182 mer) DNA. Non-Ebox DNA (with TTAGCA; 182-mer) did not induce any phase separation under the concentrations evaluated. Lower panel: DIC images of 1Ebox, 3Ebox and 7Ebox DNA only (without N-myc and Max proteins) that did not phase separate. Solution conditions were 20 mM HEPES, 150 mM KCl, 10 mM MgCl_2_, 1 mM DTT. **(D)** Quantification of dilute phase concentration of 25 µM N-myc by analytical HPLC. No phase separation was seen without 150 mM KCl (blue bar). N-myc phase separation is progressively enhanced by Max and the further addition of 1Ebox DNA (at 5 and 10 µM; green and magenta bars, respectively). Non-Ebox DNA (at 5 and 10 µM; cyan bars) did not change the N-myc dilute phase concentration and therefore did not cause phase separation. Data are mean ± SD (n = 3 samples). (**E)** Phase-separated condensates contain N-myc, Max and DNA oligonucleotides as evidenced by fluorescence imaging of LD-555-labeled N-myc, Rhodamine Red X-labeled Max and Alexa 488-labeled DNA oligonucleotides.

Phase separation of purified c-Myc-mEGFP (12 μ M) was previously reported ^23^. Here we characterized the effect of DNA on phase separation of N-myc. N-myc/Max complexes (at 10 and 3 μM, respectively) did not phase separate, but the addition of 1Ebox DNA into the N-myc/MAX solution promoted phase separation (Fig. 4C). 3Ebox and 7Ebox DNA (with 3 and 7 repeats of E-box sequences, respectively) promoted phase separation at lower concentrations, whereas the addition of non-Ebox DNA (of the same length as 7Ebox DNA), did not result in phase separation. E-box DNA in the absence of protein did not undergo phase separation (Fig. 4C).

We further quantified these effects using an analytical HPLC approach that allowed us to separate the components in the dilute phase and determine the concentration of N-myc ^54^ (Fig. 4D), thereby reporting directly on the driving force for phase separation of the system under different conditions. In a buffer without excess salt, N-myc did not undergo phase separation, and we therefore recovered the input concentration (of 25 μM). In a buffer containing 150 mM KCl, the dilute phase concentration of N-myc was ∼ 20 μM, reflecting that N-myc molecules in excess of 20 μM concentration were incorporated into condensates. The addition of MAX reduced the dilute phase concentration of N-myc further. While the addition of non-Ebox DNA did not promote stronger phase separation, the successive addition of 1Ebox DNA reduced the N-myc dilute phase concentration further in agreement with our qualitative microscopic results.

Together these results show that N-myc/MAX dimers phase separate, that interactions with E-box DNA enhance their phase separation, and that E-box DNA with multiple binding motifs can effectively scaffold N-myc/MAX phase separation in a pre-wetting transition (Fig. 4E), which has also been observed for other transcription factors ^55^.

### Recruitment of transcriptional machinery to N-myc condensates requires both TAD and bHLH-LZ domains

Because our data indicated that both the TAD and bHLH-LZ domains are important for N-myc PS, we next examined whether both domains were required for recruiting transcriptional machinery to N-myc condensates. Here we applied a chemogenetic tool named SparkDrop to drive phase separation of both N-myc mutants (Fig. 5A). SparkDrop drives protein phase separation by induction of multivalent interactions with a small molecule, lenalidomide. Briefly, SparkDrop is based on the newly engineered protein pair CEL (109 amino acids [aa]) and ZIF (31aa), which, upon addition of lenalidomide (lena), form a heterodimer (CEL⋯lena⋯ZIF). To induce PS, we fused the N-myc mutants to mEGFP and CEL (N-myc-mEGFP-CEL). To incorporate multivalency, we utilized a *de novo* designed coiled coil that is a homo-tetramer (HOTag6) ^56^. We fused ZIF, a nuclear localization signal (NLS), and a non-green fluorescent EGFP mutant (EGFP-Y66F) to HOTag6 (ZIF-NLS-EGFP(Y66F)-HOTag6). Hence, treatment with lenalidomide drives oligomerization of N-myc and induces phase separation even at concentrations that are below the saturation concentration of N-myc.

**Fig. 5.**
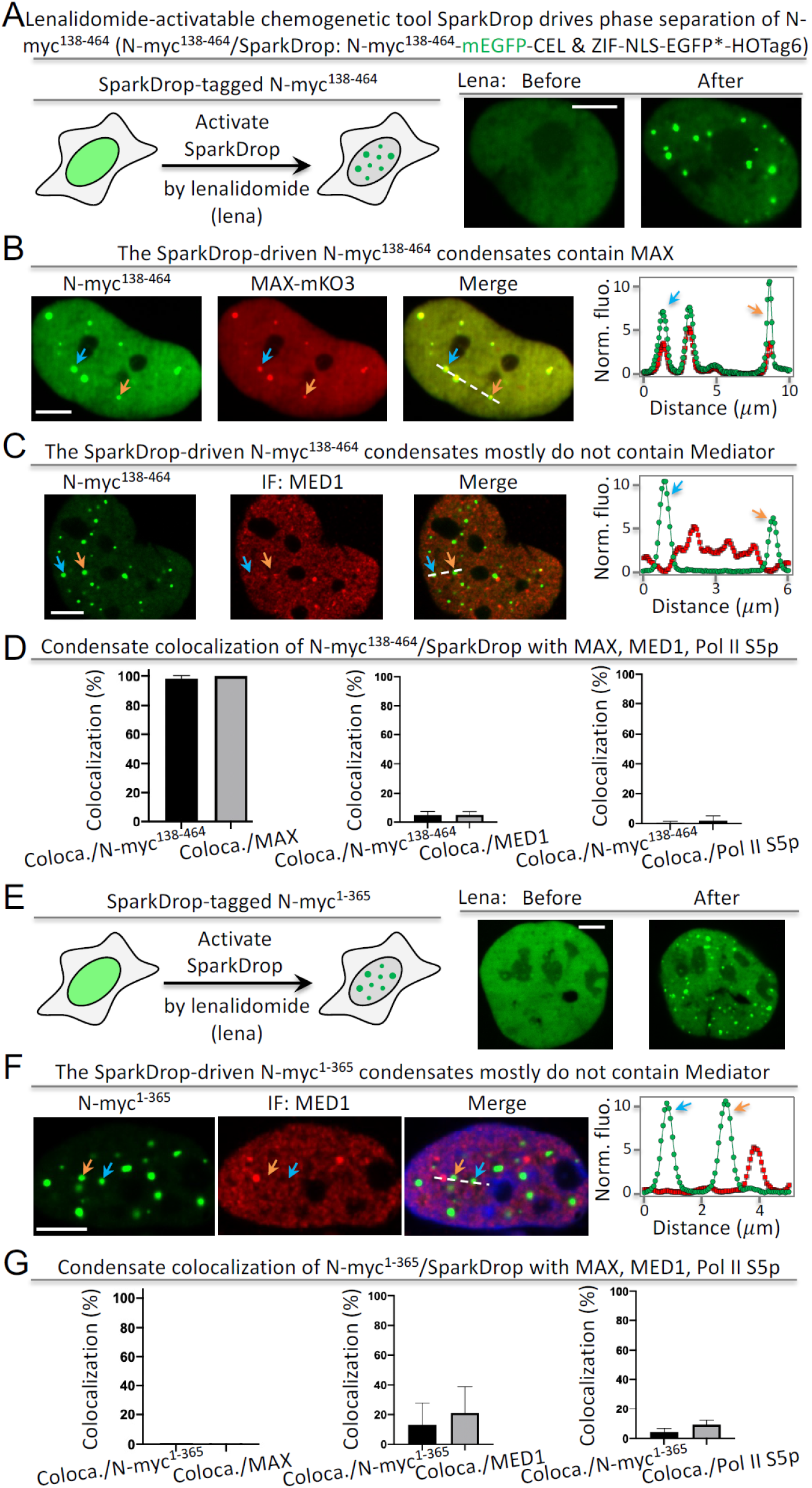
Both TAD and DNA-binding domain are required for transcriptional activity of N-myc condensates. (**A**) Lenalidomide-activatable SparkDrop drives phase separation of a truncated N-myc lacking TAD. (**B**) Fluorescence images of N-myc^138–464^/SparkDrop with MAX-mKO3. (**C**) Fluorescence images of SparkDrop-driven N-myc^138–464^ condensates and MED1. (**D**) Percentage of N-myc^138–464^/SparkDrop condensates that colocalize with puncta of other components. The percentage is determined by the ratio of coloca./N-myc^138–464^ = number of colocalized condensates between N-myc^138–464^/SparkDrop and MAX divided by number of N-myc^138–464^/SparkDrop condensates. Equivalent analysis for other pairs is also shown. Data are mean ± SD (n = 15 cells). (**E**) Lenalidomide-activatable SparkDrop drives phase separation of a truncated N-myc lacking DNA-binding domain. (**F**) Fluorescence images of SparkDrop-driven N-myc^1–365^ condensates and MED1. (**G**) Percentage of N-myc^1–365^/SparkDrop condensates that colocalize with puncta of other components. The percentage is determined by the ratio of coloca./N-myc^1–365^ = number of colocalized condensates between N-myc^1–365^/SparkDrop and MAX divided by number of N-myc^1–365^/SparkDrop condensates. Equivalent analysis for other pairs is also shown. Data are mean ± SD (n = 15 cells). Scale bars: 5 μm (A – C, E, F).

First, we demonstrated that SparkDrop induced phase separation of TAD-deleted N-myc^138–464^ upon addition of lenalidomide (Fig. 5A). The condensates recruited the DNA-binding partner MAX as expected (Fig. 5B). In contrast, most of the N-myc condensates did not contain MED1 (Fig. 5C) or Pol II S5p (Supporting Fig. S7). Most (98%) of the N-myc condensates contained MAX, whereas MED1 and Pol II S5p showed ∼5% and 0.4% colocalization, respectively (Fig. 5D). These data thus show that the TAD domain is critical for recruitment of the transcriptional machinery to N-myc condensates.

Next, we showed that SparkDrop was also able to drive phase separation of bHLH-LZ-deleted N-myc^1–365^ (Fig. 5E). The majority of these condensates contained no MED1 or Pol II S5p, with ∼13% and ∼4% colocalization, respectively (Fig. 5F, Supporting Fig. S8), indicating that they are largely inactive for gene transcription. As expected, these condensates contained no MAX (Fig. 5F). Therefore, our data show that the DNA-binding domain is also critical for assembling transcriptionally active N-myc condensates. Together, our results indicate that the TAD and the DNA-binding domains are both required to recruit the transcriptional machinery, and that condensate formation itself does not automatically result in that.

### The chemogenetic tool SparkDrop decouples N-myc PS from protein abundance

We demonstrated that SparkDrop enabled us to drive PS without changing protein levels, thus decoupling phase separation from a need to increase protein abundance. We tagged N-myc with SparkDrop (N-myc/SparkDrop). We first explored the response of N-myc/SparkDrop-expressing cells to treatment with lenalidomide. Lenalidomide activated SparkDrop and induced condensate formation within 6 – 10 minutes (Fig. 6A). The total fluorescence of N-myc showed little change during that time, suggesting that N-myc protein levels were constant in the nucleus. DMSO alone did not induce N-myc phase separation, and lenalidomide could not drive N-myc phase separation in the absence of the HOTag6. Furthermore, without N-myc, SparkDrop did not form condensates even upon addition of lenalidomide (Supporting Fig. S9). We also demonstrated that in the absence of lenalidomide, N-myc/SparkDrop undergoes PS with a saturation concentration of ∼330 – 400 nM (Supporting Fig. S10), similar to that of N-myc-mEGFP, indicating that the SparkDrop tag itself had little effect on N-myc’s phase behavior. Lastly, we showed that N-myc/SparkDrop condensates were able to fuse and coalesce, indicating that they are liquid droplets with an inverse capillary velocity of 1.31 ± 0.29 (s/μm), which is similar to that of N-myc-mEGFP (Supporting Fig. S11).

**Fig. 6.**
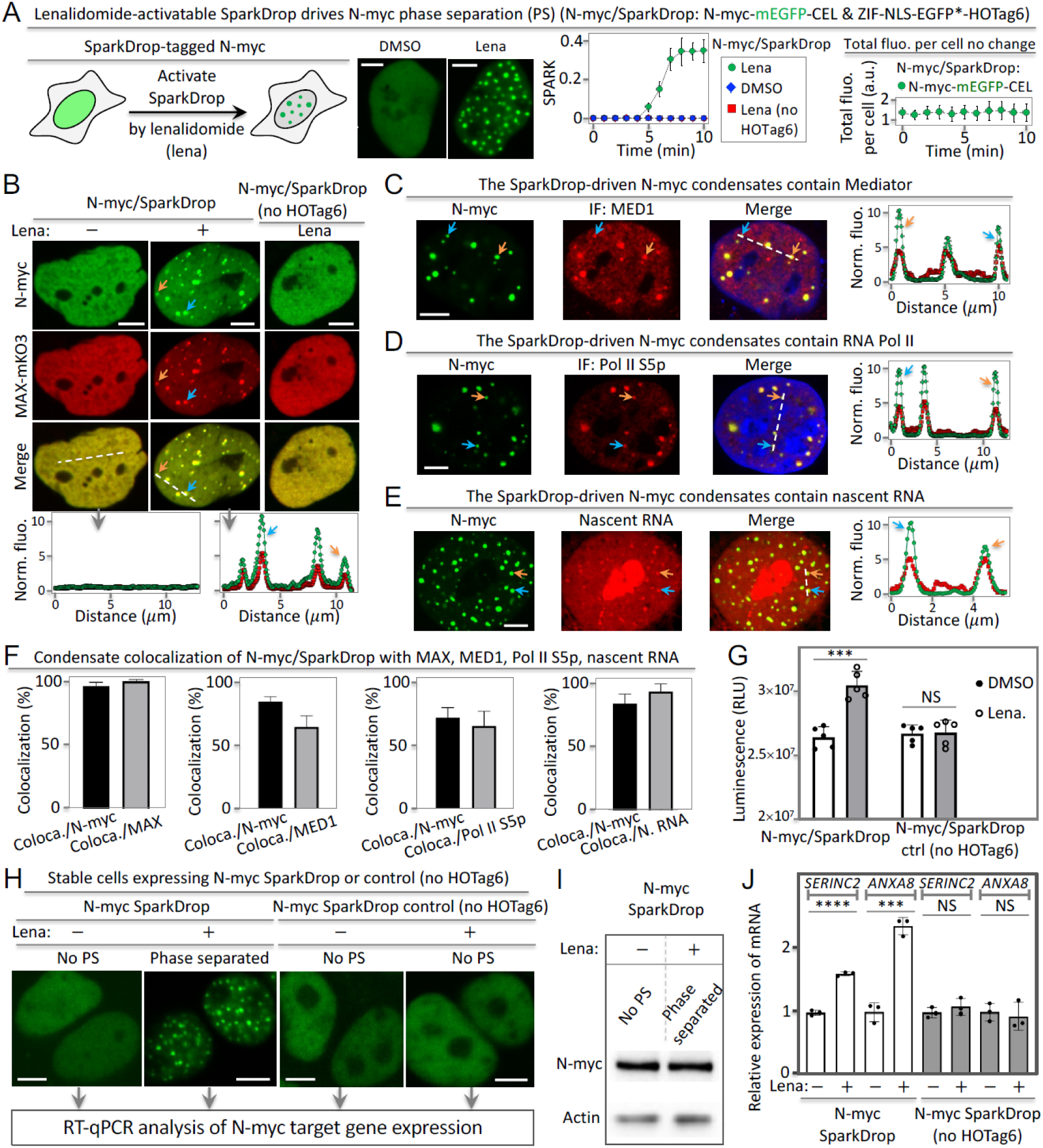
The chemogenetic tool SparkDrop reveals role of N-myc phase separation on transcription. (**A**) SparkDrop drives N-myc phase separation without change of protein level. (**B**) SparkDrop-driven N-myc condensates contain DNA-binding and dimerization partner MAX. (**C – E**) Fluorescence images showing SparkDrop-driven N-myc condensates contain transcriptional machinery including MED1 (C), RNA Pol II S5p (D), and nascent RNA (E). (**F**) Percentage of N-myc/SparkDrop condensates that colocalize with puncta of other components. The percentage is determined by the ratio of coloca./N-myc = number of colocalized condensates between N-myc/SparkDrop and MAX divided by number of N-myc/SparkDrop condensates. Equivalent analysis for other pairs is also shown including nascent RNA (N. RNA). Data are mean ± SD (n = 13 cells). (**G**) Quantitative analysis of SH-EP cell growth using CellTiter-Glo with N-myc in non-phase-separated vs phase-separated state. Luminescence was measured after the cells were treated with DMSO or lenalidomide (1μM) for 72 hrs. Data are mean ± SD (n = 5). *** P-value < 0.001. (**H**) Fluorescent images of stable cells expressing SparkDrop-tagged N-myc or the control. The cells were treated with lenalidomide or DMSO, followed by RT-qPCR analysis. No PS: no phase separation. (**I**) Western blot showing N-myc protein abundance level. (**J**) RT-qPCR analysis of the expression levels of two N-myc-regulated genes in cells without and with phase separation of N-myc. Data are mean ± SD (n = 3). ****P-value < 0.0001. *** P-value < 0.001. NS, not significant. Scale bars: 5 μm (A – E, H).

Next, we tested whether N-myc/SparkDrop condensates bear the hallmarks of transcriptional activity. First, the N-myc/SparkDrop condensates contained the DNA-binding and dimerization partner MAX, with a colocalization of ∼ 95% (Fig. 6B). Second, the N-myc/SparkDrop condensates contained transcriptional machinery including MED1 and Pol II S5p (Fig. 6C, D). Lastly, the N-myc/SparkDrop condensates contained nascent RNA (Fig. 6F). Colocalization of MAX, MED1, Pol II S5p, and nascent RNA with N-myc condensates was 95%, 85%, 72%, and 80%, respectively (Fig. 6F). These data thus show that SparkDrop is an appropriate tool to form condensates with the hallmarks of those formed by N-myc but can do so without changing the protein concentration. Therefore, it allows us to decouple the effect of phase separation on N-myc-mediated transcription and function from the effect of increasing N-myc levels.

As a first step in this direction, we tested whether N-myc phase separation had an effect on cell survival. The cell viability was increased by 15 ± 4% when N-myc/SparkDrop expressing cells were treated with lenalidomide as compared to the DMSO-treated cells (Fig. 6G). In cells expressing the N-myc/SparkDrop control without the HOTag6, lenalidomide had no effect on cell viability. Therefore, phase separation of N-myc leads to a significant survival advantage for cells, even at the same N-myc concentration (Fig. 6A, H, I).

### PS of N-myc regulates gene transcription

Next, we examined whether lenalidomide/SparkDrop-induced phase separation of N-myc regulated gene transcription. We chose two key N-myc-regulated genes which are known for their oncogenic character in promoting cell proliferation and survival, serine incorporator 2 (*SERINC2*) and annexin A8 (*ANXA8*) ^57,58^, and examined whether their transcription was altered upon N-myc phase separation. RT-qPCR analysis revealed that the mRNA levels of *SERINC2* and *ANXA8* were significantly higher for cells with N-myc condensates than for cells without condensates (Fig. 6J). Lenalidomide treatment of cells expressing the N-myc/SparkDrop control without HOTag6 did not result in increased transcription. Therefore, our data indicate that the process of N-myc phase separation enhances transcriptional activity of key proliferative genes, in agreement with our results that phase separation also promotes survival of SH-EP cells.

### Phase-separated N-myc regulates the transcriptome

We next tested the effect of phase-separated N-myc (induced by lenalidomide/SparkDrop) on the transcriptome without changing N-myc abundance in the nucleus (Fig. 7A). We treated the engineered cells expressing N-myc/SparkDrop with and without lenalidomide, followed by RNA-seq.

**Fig. 7.**
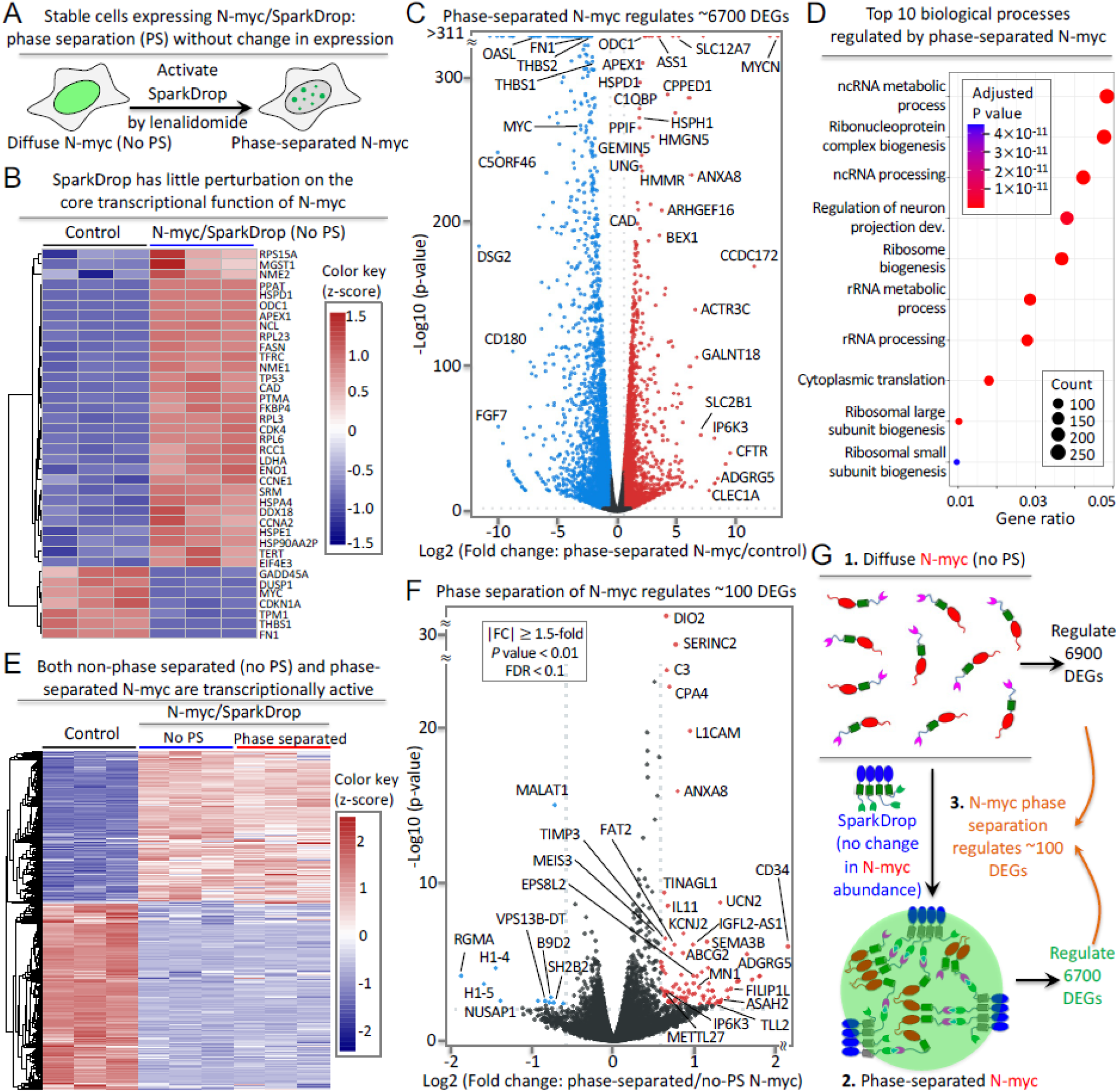
Phase separation of N-myc regulates transcription of a small percentage of genes. (**A**) Schematic of SparkDrop-based N-myc phase separation without change of protein abundance levels. (**B**) Heatmap of gene regulation in response to N-myc/SparkDrop-expression (without phase separation) on core target genes of N-myc. (**C**) Volcano plot showing DEGs that are regulated by phase-separated N-myc compared to no N-myc expression. The mRNAs with significant up- and down-regulation (|FC| ≥ 1.5, p-value < 0.01, FDR < 0.1) are marked in red and blue, respectively. Black dots represent mRNAs with no significant changes. (**D**) GO enrichment analysis of biological processes that are regulated by phase-separated N-myc. (**E**) Heat map showing DEGs regulated by diffuse N-myc (non-phase separated) and phase-separated N-myc. The color represents z-scores. (**F**) Volcano plot showing DEGs that are regulated by phase separation of N-myc (i.e. phase-separated N-myc compared to non-phase separated N-myc). The mRNAs with significant up- and down-regulation (|FC| ≥ 1.5, p-value < 0.01, FDR < 0.1) are marked in red and blue, respectively. Black dots represent mRNAs with no significant changes. (**G**) Schematic showing transcriptional regulation by diffuse N-myc (no phase separation), phase separated-N-myc, as well as N-myc phase separation, i.e., the difference between them.

First, we tested whether the SparkDrop tag itself perturbed the core transcriptional function of N-myc. We examined a previously established list of 41 core Myc-dependent genes ^59^, and found that 38 out of 41 were expressed upon N-Myc/SparkDrop expression in SH-EP cells in the absence of lenalidomide while they were not expressed in control SH-EP cells that do not express N-myc (Fig.7B). This suggested that the SparkDrop system caused little perturbation of the core transcriptional function of N-myc and was thus an appropriate tool to use.

Second, we tested the effect of SparkDrop-tagged N-myc in the absence of lenalidomide on the transcriptome. We calculated differentially expressed genes (DEGs; p-value < 0.01, |Log_2_FC| ≥ 0.58, FDR < 0.1), which revealed a change of gene expression by N-myc/SparkDrop. We identified ∼6900 DEGs with ≥ 1.5-fold change in transcript level, including ∼3100 up-regulated genes and ∼3800 down-regulated genes (Supporting Fig. S12, Supporting Excel File 1), consistent with previous reports of the effect of N-myc-expression on gene transcription ^60,61^. Gene ontology (GO) enrichment analysis reveals that the DEGs are strongly linked to several N-myc-regulated biological processes including ribosome biogenesis (Supporting Fig. S13). Therefore, our results indicate that SparkDrop has no or little perturbation on the N-myc transcriptional function and that N-myc activates known transcriptional programs also in SH-EP cells.

Third, we tested whether phase-separated N-myc induced by lenalidomide/SparkDrop is functional and transcriptionally active in regulating the core N-myc target genes and the transcriptome. We characterized transcriptomic changes upon lenalidomide treatment of N-myc/SparkDrop expressing cells (relative to the control cells without N-myc/SparkDrop). ∼6700 DEGs (p-value < 0.01, |Log_2_FC| ≥ 0.58, FDR < 0.1) were identified with ≥ 1.5-fold change in gene expression, including up-regulation of 3025 genes and down-regulation of 3636 genes (Fig. 7C, Supporting Excel File 2). This result is consistent with our imaging data that the N-myc/SparkDrop condensates contain transcriptional machinery (Fig. 6B – E). GO enrichment analysis reveals that the phase separated-N-myc regulates several known biological processes, including ribosome biogenesis (Fig. 7D) ^60,61^.

### Phase separation of N-myc differentially modulates the transcriptome

We then wondered to which extent transcriptional changes are different between N-myc/SparkDrop diffuse expression (without lenalidomide treatment and thus no phase separation) and phase-separated N-myc/SparkDrop (with lenalidomide treatment), i.e. how much phase separation contributed to function on top of expression of diffuse N-myc. Of note, these conditions have the same expression levels but the former has no N-myc phase separation and thus no condensates while the latter has N-myc phase separation and thus contains N-myc condensates. We determined DEGs (p-value < 0.01, |Log_2_FC| ≥ 0.58, FDR < 0.1) by comparing the RNA-seq data of the engineered cells expressing the N-myc/SparkDrop with and without lenalidomide. We identified 99 DEGs that were differentially expressed with ≥ 1.5-fold change in transcript levels, including 88 up-regulated genes and 11 down-regulated genes (Fig. 7E, F, Supporting Excel File 3). These genes are therefore regulated differentially by the process of N-myc phase separation. In contrast, lenalidomide treatment in engineered cells expressing the N-myc/SparkDrop control without HOTag6 had little effect on transcriptional activity; only 10 DEGs were identified and none of them overlapped with the N-myc PS-regulated genes when we compared the RNA-seq data of lenalidomide versus DMSO-treated cells (Supporting Excel File 4).

Which genes are affected by phase separation? Several genes that are up-regulated by N-myc expression, for example, *C3* and *SERINC2*, are further up-regulated by phase separation. This means that: 1) *SERINC2* and *C3* are up-regulated by diffuse N-myc (compared to the control); 2) and their transcription is further increased when N-myc undergoes phase separation forming condensates. This is also in agreement with our qRT-PCR results for *SERINC2* (Fig. 6J). In total, < 3% of the up-regulated genes are further enhanced by N-myc phase separation. Among the genes that are down-regulated by N-myc expression, several are further down-regulated by phase separation, including *RGMA* and *SH2B2*. In total, phase separation suppresses transcription of < 1% N-myc-repressed genes. The corollary of these findings is that transcription of most genes is not influenced by N-myc phase separation. In other words, formation of N-myc condensates from the diffusively distributed state does not affect transcript levels for the majority of the genes. Therefore, N-myc phase separation has no function on transcription of these genes, even though the N-myc condensates are transcriptionally active.

Importantly, many of the genes up-regulated by N-myc PS are oncogenes. For example, the complement component *C3* promotes cancer progression ^62,63^. *SERINC2* promotes cell proliferation and survival^58^. On the other hand, many of the genes down-regulated by N-myc PS are tumor suppressors. For example, *RGMA* is frequently inactivated and down-regulated in most colorectal cancers ^64^. *MALAT1* suppresses metastasis as its knockout induces metastatic ability of human breast cancer cells ^65^. Down-regulation of *MALAT1* has also been shown in Myc-driven tumor samples ^60^. In conclusion, N-myc phase separation alters the expression of a small subset of Myc-responsive genes, but these are enriched with oncogenes and tumor suppressors that have previously been linked to oncogenesis.

## Discussion

In this work, we developed and used a powerful chemogenetic tool, SparkDrop, that allows us to induce phase separation without changing expression levels of the driver of phase separation, here N-myc. Treatment with lenalidomide to activate SparkDrop and induce phase separation of N-myc in the neuroblastoma SH-EP cell line left N-myc levels in the nucleus unchanged but resulted in rapid formation of condensates on a timescale of 10 minutes. These condensates appeared at genomic loci of Myc-responsive genes, recruited MAX, transcriptional machinery and contained nascent RNA; hence, they have the hallmarks of active transcriptional condensates. Our transcriptome analysis further confirmed that both the diffuse expression and phase separation of N-myc resulted in the transcriptional regulation of core Myc-responsive genes. However, we also found in our comparison of the phase-separated vs the diffuse state of N-myc, a unique comparison enabled by SparkDrop, that most Myc-responsive genes do not experience expression changes upon N-myc phase separation. Only ∼3% of genes that are upregulated by Myc expression show further upregulation upon phase separation. <1% of Myc-downregulated genes show further down-regulation upon phase separation. Our analysis shows that these DEGs are enriched with oncogenes and tumor suppressors, respectively, and that this mechanism may therefore contribute to oncogenesis. However, our results also emphasize that diffuse complexes formed by N-myc with the transcriptional machinery are sufficient for strong transcriptional activity and that phase separation is a by-product of high N-myc expression rather than a state that alters function broadly.

Phase separation has been increasingly linked to transcription, and there is growing evidence that transcriptional condensates occur in functional and developmental states ^66^ as well as in disease ^29,32,37,67,68^. But whether phase separation mediates unique functions in transcription that are not mediated by diffuse complexes has remained controversial. Here we show that triggering of phase separation influences the expression of only a small fraction of N-myc-responsive genes, cautioning the assignment of special functions to condensates without proof. Our results demonstrate the transcriptional activity of N-myc condensates; they recruit transcriptional machinery, appear at target genes and contain nascent RNA. Furthermore, the transcriptional activity across most genes remains constant upon phase separation, showing that they represent a state with similar activity as the diffuse state, at the same total N-Myc concentration. The formation of condensates due to an increase in protein concentration would therefore likely result in increased function, but our results lead us to predict that the same increase in protein concentration without phase separation would result in a similar functional increase. Biomolecular condensates can possess emergent properties that range from activating reactions ^69^ to filtering noise ^70,71^ and changing translation patterns ^72^, but to elucidate these functions requires careful characterization and dissociation of function ^73^.

Why is the transcription of a fraction of genes altered upon N-myc phase separation? These effects may be related to emergent properties of condensates such as their ability to coalesce. This is a property not inherent in small complexes, which have a fixed size distribution at a given concentration and do not grow over time ^34^. Coalescence of transcriptional condensates can alter chromatin structure and thereby likely bring genes in the vicinity of strong enhancers that alter their expression ^32^. Such effects may become more dominant the longer TF levels are elevated, although our data show that N-myc condensates disassemble during mitosis and that their lifetimes and their ability to coalesce and ripen are therefore naturally limited.

We conclude that N-myc condensates do not mediate a super-proportional fraction of the Myc-related activity. Condensates may nonetheless represent interesting drug targets because therapeutics may be enriched within them not only by direct binding of targets but also through binding of other components in condensates and solubility effects ^37,38^. Whether such a therapeutic strategy would be successful likely depends on the fraction of N-myc within condensates. This is ∼30-50% in our experiments and may be a higher fraction in cancers with high overexpression of Myc due to copy number changes, chromosomal translocations, or upstream oncogenic signaling^3–5^.

While our work addresses the role of phase separation specifically in N-myc transcriptional activity, we expect that many of our findings will hold in other systems and will thus open new directions for understanding the role of phase separation in transcription in general.

## Acknowledgments

We thank Hiten Madhani, Bo Huang, Vijay Ramani, and Eric Holland for critical suggestions.

## Author contributions

X.S. conceived the project. J.Y. performed N-myc phase separation and colocalization with other proteins in cells. C-I.C. conducted imaging of small molecule-induced N-myc phase separation and analyzed colocalization with other proteins. C-I.C. performed and analyzed nascent RNA labeling, RT-qPCR and RNA-seq. J.Y., J.K. and W.A.W planned and performed experiments to analyze expression of endogenous N-myc protein in the neuroblastoma cells. H.L. processed RNA-seq data. H.L., C-I.C, J.Y., Q.Z., X.Y., X.S., Y.S. analyzed RNA-seq data. A.N. and T.M. designed and analyzed the in vitro experiments. A.N. conducted the in vitro experiments. J.Y., C-I.C, T.M., X.S. wrote the manuscript. All authors contributed to the final draft.

## Data and materials availability

All data are available in the main text or the supplementary materials.

## Supplementary Materials

Supporting excel files 1 – 4 (list of DEGs from RNA-seq)

## Supplementary Information

### Materials and Methods

#### Plasmid construction

N-myc and its truncations were amplified from cDNA library. MAX was amplified from The CCSB Human ORFeome Collection (donated by Marc Vidal). mKO2-N1 was a gift from Michael Davidson & Atsushi Miyawaki (Addgene plasmid # 54625) and mKO3 was obtained by introducing the M176F mutation to mKO2 for brighter fluorescence in cells (Mastop et al., 2017; Tsutsui et al., 2008). N-myc-mEGFP, N-myc^1–365^–mEGFP, N-myc^138–464^–mEGFP, MAX-mKO3 were created by linking the DNA sequence of N-myc and its truncations to mEGFP, MAX to mKO3 with a 10-AA GSSGGSGGGT linker. To create N-myc-mEGFP-CEL fusions, CEL was amplified from full length CRBN and cloned into N-myc-mEGFP plasmid, with a 10-AA linker between mEGFP and CEL. The full-length N-myc was replaced with its truncations resulting in N-myc^1–365^–mEGFP-CEL and N-myc^138–464^–mEGFP-CEL. All the sequences were cloned into pHR_SFFV (Addgene plasmid #79121) vector by standard restriction enzyme digestion and ligation method and confirmed by exhaustively sequencing. All constructs in this study are listed in Table S1.

#### Cell culture

SH-EP (University of California, San Francisco, Cell and Genome Engineering Core) cells were cultured at 37 °C and 5% CO_2_ in Dulbecco’s Modified Eagle medium (DMEM) high glucose, supplemented with 10% Fetal Bovine Serum (FBS) heat inactivated, penicillin (100 units/mL) and streptomycin (100 μg/mL). Kelly (Sigma-Aldridge), CHP134 and CLB-GA (Childhood Cancer Repository, Children’s Oncology Group resource laboratory) cells were cultured at 37 °C and 5% CO_2_ in Roswell Park Memorial Institute (RPMI) 1640 Medium, supplemented with 10% Fetal Bovine Serum (FBS) heat inactivated All culture supplies were obtained from Gibco.

#### Cell viability measurement

First, SH-EP cell single clones stably expressing N-myc/SparkDrop or N-myc/SparkDrop control was generated by single-cell dilution and selection after infection of lentivirus containing N-myc/SparkDrop or N-myc/SparkDrop control. Second, cells were grown on clear plastic 96-well plates for 24 hours. Then, the cells were treated with DMSO or lenalidomide (1 µM; TargetMol, T1642) for 72 hours. CellTiter-Glo reagent (Promega G9243) was added, and the cells were gently shaken for 10 minutes. The mix content was then moved to white 96-well plates, and luminescence was measured using a Tecan Spark plate reader. For each biological repeat, 6 technical repeats (6 wells) were measured.

#### Lentivirus preparation

N-myc-mEGFP, N-myc (full-length or truncated) /SparkDrop, N-myc/SparkDrop control (no HOTag6), mKO3-MAX, H2B-mIFP, H2B-mCherry and mEGFP lentiviral plasmids were co-transfected with pAX2 and pVSVG at 3:2:1 ratio into HEK293T cells using polyethylenimine (MilliporeSigma 764965). Lentiviruses were harvested after 48 hours.

#### Confocal microscopy

Samples were imaged on a Nikon Eclipse Ti inverted microscope equipped with a Yokogawa CSU-W1 confocal scanner unit (Andor), a digital CMOS camera ORCA-Flash4.0 (Hamamatsu), a ASI MS-2000 XYZ automated stage (Applied Scientific Instrumentation) and Nikon Plan Apo λ 20X air (N.A. 0.75), Nikon Apo TIRF 60X oil (N.A. 1.49) and CFI Plan Apo λ 100X oil (N.A. 1.45) objectives. Laser inputs were provided by an Integrated Laser Engine (Spectral Applied Research) equipped with laser lines (Coherent) 405 nm for Hoechst imaging, 488 nm for GFP imaging, 561 nm for mCherry imaging and 640 nm for mIFP imaging. The confocal scanning unit was equipped with the following emission filters: 460/50 nm for Hoechst imaging, 525/50-nm for GFP imaging, 610/60-nm for mCherry imaging and 732/68 nm for mIFP imaging. Image acquisition was controlled by the NIS-Elements Ar Microscope Imaging Software (Nikon). Images were processed using NIS-Elements and ImageJ (NIH).

#### Live cell imaging

SH-EP cells were grown in glass-bottom 8-well chambers (Thermo Scientific 155411). Lentiviral infection was performed when cells were cultured to ∼70% confluence. Cells were imaged 24 hours after infection. Time-lapse imaging was performed with the aid of an environmental control unit incubation chamber (InVivo Scientific), which was maintained at 37 °C and 5% CO_2_. For imaging mitosis, 100 μM ascorbic acid was added to increase mitotic index. Lenalidomide (Ark Pharma, AK-47482) were carefully added to the cells in the incubation chamber when the time-lapse imaging was started. Image acquisition was controlled by the NIS-Elements Ar Microscope Imaging Software (Nikon). Images were processed using NIS-Elements and ImageJ (NIH). Chromosome volume was calculated based on the near infrared fluorescence from H2B-mIFP. The fusion events of the N-myc condensates and N-myc/SparkDrop condensates were analyzed by calculating the aspect ratio of the fusing condensates over time using imageJ.

#### mEGFP concentration and fluorescence intensity standard curve

We purified mEGFP protein and measured its 488 nm absorbance using a Thermo scientific nanodrop 2000C to estimate the concentration through dividing the absorbance value by extinction co-efficiency. The protein was then serial diluted between 0.037 μM and 2.35 μM, added into 8-well chambers and imaged under the same Nikon Eclipse Ti inverted microscope and parameters as for N-myc-mEGFP phase diagrams. The laser power was also measured by a Thorlabs PM100D optical power and energy meter to make sure it is stable and the same as for phase diagrams. Fluorescence intensity (counts/pixel) under 488 channel was recorded to plot the mEGFP concentration-fluorescence standard curve.

#### Colocalization analysis

We quantified the percentage of condensate colocalization for full length or truncated N-myc/SparkDrop condensates with MAX, MED1 antibody staining, RNA polymerase II S5P antibody staining, nascent RNA labeling, in single nucleus via 3D imaging using spinning disc confocal microscope at 100X. Each nucleus was sectioned into multiple slices with 0.5 μm interval. To calculate condensate number, the images were imported into ImageJ and the condensate were selected by threshold, which were then added to ROI manager. Total number of condensates was summarized from each slice. The overlapped condensates were selected using AND in ROI manager and the number of overlapped condensates was calculated using Analyze particles.

#### Phase diagrams and droplet size and amount analysis

SH-EP cells expressing N-myc-mEGFP or N-myc/SparkDrop were stained with 1 μg/mL Hoechst 33342 for 10 minutes. The cells were imaged under a spinning disc confocal microscope with 100X objective. Each nucleus was sectioned into multiple slices with 0.5 μm interval. Using the 3D Objects Counter function in ImageJ, a low threshold of fluorescence intensity was set for each cell, to choose the whole nucleus. The mean green fluorescence intensity in each nucleus was calculated by the ratio of total fluorescence intensity over volume of the nucleus. The calculated mean fluorescence intensity was used to determine the average N-myc protein concentration by comparing with the purified mEGFP concentration vs fluorescence intensity. Next, a higher threshold was set and adjusted for each cell to select the N-myc droplets. Total droplet fluorescence intensity in each nucleus was calculated using ImageJ. SPARK value was then determined and plotted to generate the phase diagram.

#### Sequence of N-myc, Max constructs and DNA oligonucleotides

Labeled (Alexa488) and unlabeled DNA oligonucleotides were synthesized by Integrated DNA Technologies, Inc. (IDT, USA). The sequences of proteins and DNA oligonucleotides used in the current study are provided in the table below:

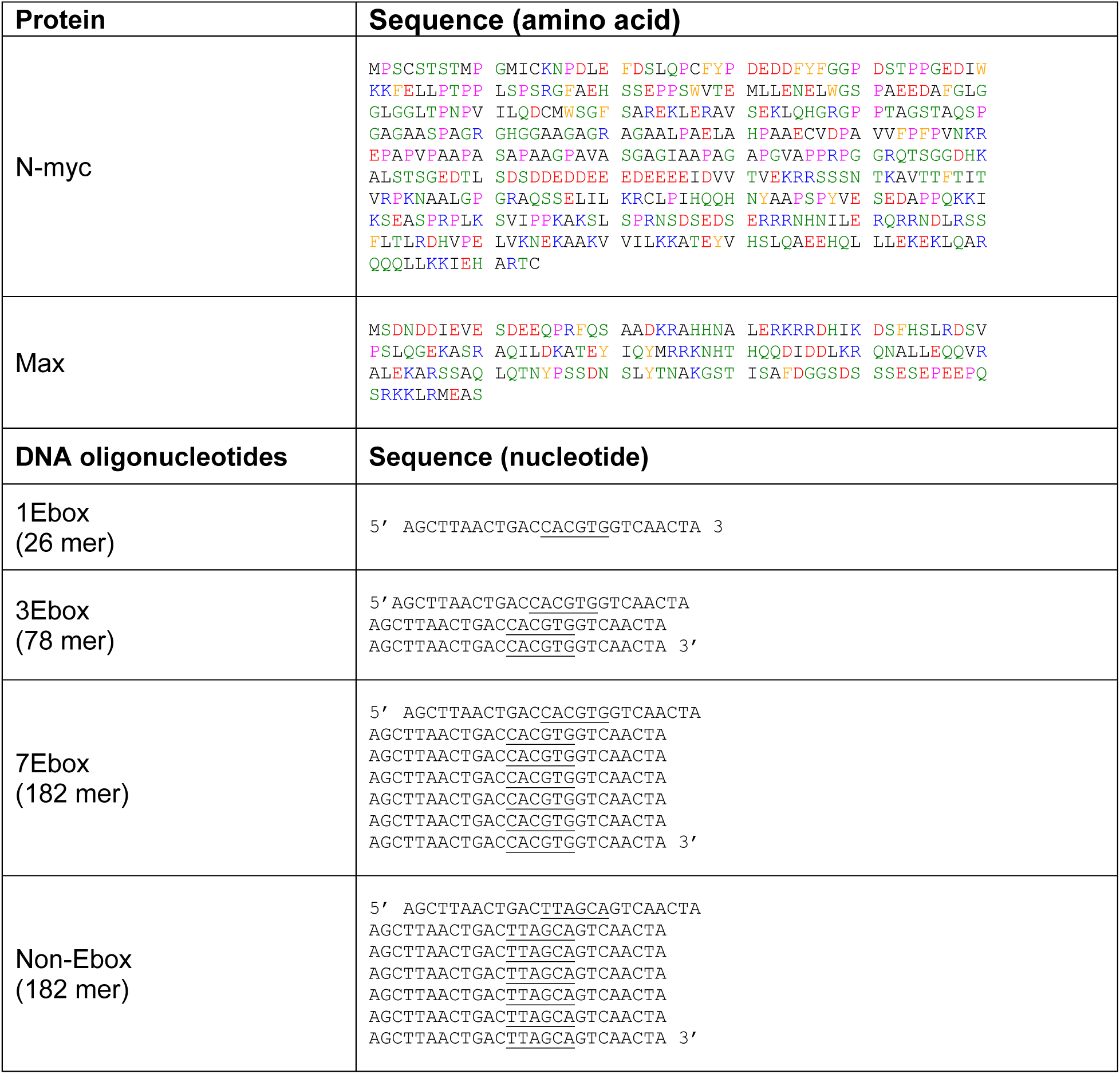

#### Expression and purification of N-myc & Max proteins

The sequences for human N-myc and Max proteins were synthesized (Thermo Fisher Gateway cloning) along with a coding sequence for an N-terminal TEV protease cleavage site (ENLYFQGS) including 5’ and 3’ attB sites for cloning into expression vectors. The sequences were codon optimized for expression in an *E. coli* system. The sequences from the pDONR221 vector were transferred using Gateway™ LR Clonase™ Enzyme mix (Invitrogen, USA) to the expression vector pDEST17 (Thermo Fisher, USA), containing N-terminal 6xHis-tag sequence. Both proteins were expressed in *E. coli* strains [BL21(DE3) pLysS for Myc and BL21-Gold(DE3) for Max]. Cultures were grown to an OD_600_ = 0.8 and induced with 1 mM IPTG followed by overnight expression at 18 ℃. Pelleted cells were resuspended in 50 mM Tris pH 7.5, 1M NaCl, 30 mM Imidazole and 5 mM β-mercaptoethanol. Further, the pellets were lysed using sonication on ice. Cell lysates were centrifuged and insoluble inclusion bodies were collected. The inclusion bodies were solubilized in a buffer containing 20 mM Tris pH 7.5, 6 M GdmHCl, 50 mM NaCl, 50 mM Imidazole and 5 mM β-mercaptoethanol overnight at 4°C.

Solubilized inclusion bodies were cleared by centrifugation at 16,000 rpm and supernatants were loaded onto columns of Chelating Sepharose Fast Flow^TM^ resin (GE Healthcare, USA) charged with nickel sulfate. The columns were washed with 3 column volumes of 20 mM Tris pH 7.5, 4 M urea, 50 mM imidazole. Protein elution from the Ni-NTA resin was done using 4 M urea, 20 mM Tris pH 7.5, 500 mM imidazole. The eluted proteins were used to cleave the 6xHis-tag by TEV cleavage in 20 mM Tris pH 7.5, 2 M urea, 50 mM NaCl, 0.5 mM EDTA, 1 mM DTT buffer, overnight at 4°C. The cleaved proteins were passed over Ni-NTA columns and the flowthrough and wash fractions were collected. The proteins were concentrated using Amicon centrifugal filters (3,000 MWCO). The proteins were further purified using a S75 Superdex size exclusion column (GE Healthcare, USA) with buffer containing 20 mM Tris pH 7.5, 2 M GdmHCl, 50 mM NaCl and 1 mM TCEP. The identity of the proteins was confirmed using intact mass spectrometry. N-myc and Max were stored in denaturing buffer with 4 M GdmHCl, 20 mM Tris pH 7.5, 50 mM NaCl and 1 mM TCEP at 4°C. The molar absorptivity of N-myc and Max were taken as 29520 M^−1^ cm^−1^ and 5960 M^−1^ cm^−1^, respectively.

#### Refolding of Myc and Max proteins

Both N-myc and Max proteins were individually refolded using serial steps of dialysis as previously reported (Farina et al., 2004). The proteins were dialyzed in Spectra/Por 3 (3.5 kD MWCO) Dialysis Tubing (Fisher Scientific, USA). Buffer containing 20 mM HEPES pH 7.5, 50 mM KCl, 10 mM MgCl_2_ and 1 mM DTT with decreasing levels of urea was used for dialysis during each refolding step (4 M urea, 2 M urea, 1 M urea; each step for 1h) at 4°C. In the final step, buffer with 20 mM HEPES pH 7.5, 50 mM KCl, 10 mM MgCl_2_ and 1 mM DTT was used for overnight dialysis at 4°C. The refolded N-Myc and Max proteins were used in a 3:1 ratio for all assays. The proteins were freshly refolded for each experiment.

#### Far UV CD spectroscopy

To confirm refolding, samples were dialyzed in 20 mM phosphate buffer pH 7.5, 50 mM KCl, 10 mM MgCl_2_ and 1 mM DTT. The CD spectra were measured for N-myc (10 µM) and Max (3 µM) individually and as a complex in presence of 1Ebox DNA (50 nM). The spectra were recorded with a J-1500 spectrophotometer (Jasco, USA) at 25 ℃. Samples were measured in a 0.1 mm pathlength cuvette (Hellma, Germany). The spectra were accumulated thrice with a response time of 4 s, 1 nm data pitch and 1 nm bandwidth from 195 to 250 nm.

#### Fluorescence anisotropy binding assay

A SpectraMax^TM^ fluorescence polarization plate reader was used to assay the binding of Alexa488-labeled 1 Ebox DNA oligonucleotide to a mixture of N-myc / Max protein. The assays were performed using 20 mM HEPES, 150 mM KCl, 10 mM MgCl_2_, 1 mM DTT, 0.01% NP-40 in a non-binding flat-bottom 384-well black plate. The concentration of fluorophore-labeled DNA was fixed at 50 nM and proteins are titrated in the ratio of Myc / Max at 3:1. The dissociation constant (*K_d_*) of the interaction between proteins and DNA was estimated by fitting the experimentally acquired anisotropy values in Origin Pro 8.0 (Roehrl et al., 2004).

#### Microscopy of N-myc, Max and DNA

For assessing the driving force for phase separation, samples of N-myc (10 µM), Max (3 µM) and DNA (titration from 0.25 -10 µM for 1Ebox, 3Ebox and 7Ebox) were incubated in buffer containing 20 mM HEPES pH 7.5, 150 mM KCl, 10 mM MgCl_2_ and 1 mM DTT at 25 ℃. 4 µL of each sample was sandwiched between an alcohol-cleaned glass slide and coverslip using a 3M 300 LSE high-temperature double-sided tape (0.34 mm) with a cutout window. The microscopy images were captured using a Nikon Eclipse Ni-E Widefield microscope with a 20X objective. For fluorescence imaging, 1% of the labeled component (LD655-labeled N-myc, Rhodamine red X-labeled Max and Alexa488-labeled DNA oligos) were added to the samples and imaged using the fluorescence Andor camera on the Nikon Eclipse Ni-E Widefield microscope with a 20X objective. For each sample, at least two replicates were imaged.

#### HPLC phase separation assay

The N-myc concentration in the dilute phase in the presence of Max and/or DNA oligonucleotides was determined by using analytical HPLC as per the previously established protocol (Bremer et al., 2022). The input concentration of N-myc was 25 µM for all samples, which were incubated in the presence (or absence) of Max (8 µM) and DNA oligonucleotides (5 µM and 10 µM with 1Ebox or Non-Ebox sequence) in 20 mM HEPES pH 7.5, 150 mM KCl, 10 mM MgCl_2_ and 1 mM DTT. N-myc protein without 150 mM KCl was included as a control (which did not phase separate). All solutions were incubated at 25℃ for 20 min, then centrifuged for 15 min at 12,000 rpm to separate the dilute and dense phases. An equivalent amount of dilute phase was removed from all tubes and diluted as needed by a defined value. The dilute phase solution was then applied to the HPLC C18 (ReproSil Gold 200; Dr. Maisch) reverse-phase column to determine the concentration of N-myc (at 280 nm wavelength). A buffer system with a gradient of H_2_O + 0.1% TFA (trifluoroacetic acid, Sigma-Aldrich, USA) with acetonitrile (Alfa Aesar, USA) was used as a solvent system. A standard curve with known concentrations of purified N-myc protein was subjected to HPLC similarly to determine the concentrations of N-myc in the dilute phase for each of the conditions.

#### Immunofluorescence

SH-EP cells were cultured in a glass-like-bottom 96 well plate, infected with N-myc-mEGFP (full-length or truncated) or N-myc/SparkDrop viruses, and fixed with 4 % paraformaldehyde in PBS at 24 hours post-infection. SH-EP cells were permeabilized with PBST (0.1% Triton X-100 in PBS) and blocked with 2% BSA and 10% goat serum in PBS for 30 minutes. The cells were next incubated overnight with primary antibodies in blocking buffer. After washing three times with PBST, the cells were incubated with secondary antibodies at room temperature for 1 hour. After washing three times with PBST, 100 μL of PBS + 1 μg/mL Hoechst 33342 was added to each well and the cells were imaged after 10 minutes. Kelly, CHP134, SH-EP and CLB-GA cells were grown on glass coverslips in 6-well plates for 24 hours. The cells were fixed with 4 % paraformaldehyde in PBS, permeabilized with PBST (0.1% Triton X-100 in PBS) and blocked with 5% normal goat serum in PBST for 30 minutes. The cells were incubated overnight with anti-MYCN antibody in blocking buffer at 4 °C. After washing three times with PBST, the cells were incubated with secondary antibodies at room temperature for 1 hour, then mounted onto glass slides using mounting media with DAPI (Vectashield H-1800).

Detailed antibody information, incubation time and temperature are listed below:

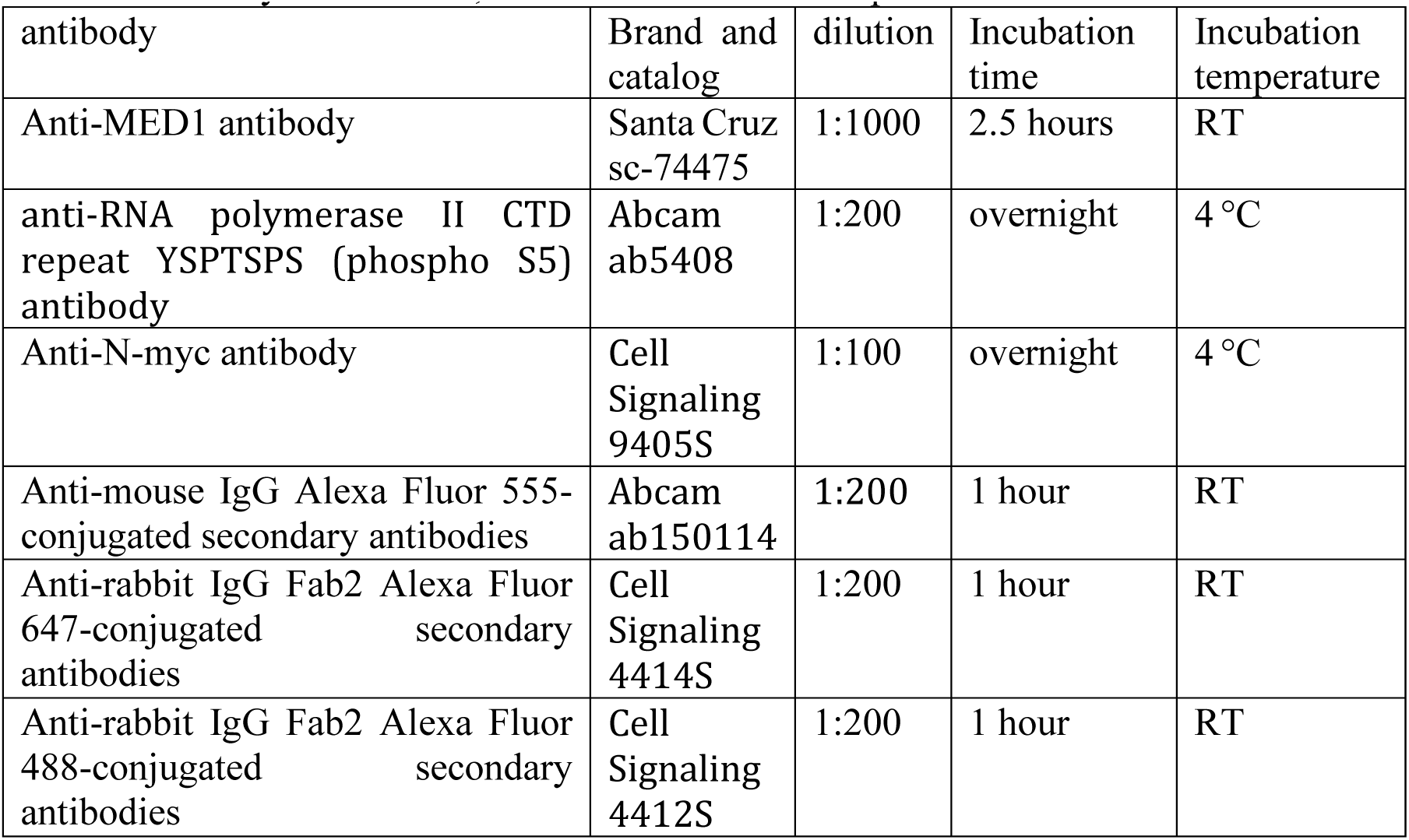

#### DNA FISH

SH-EP cells were cultured in a glass-like-bottom 96 well plate, infected with N-myc-mEGFP virus, and imaged for green fluorescence. The cells were then fixed in 3:1 methanol/acetic acid and dried, immersed in 2x Saline Sodium Citrate (SSC) (PH 7.0) for 2 minutes at room temperature (RT), dehydrated in an ethanol series (70%, 85% and 100%), each for 2 minutes at RT and dried. TP53 FISH probes (Cytocell LPH571-A) and the plate were pre-warmed at 37°C for 5 minutes. 50 μL probes were added to the well and denatured at 75°C for 2 minutes, followed by hybridization at 37°C overnight. PBS was added to surrounding wells to ensure a humid environment. After hybridization, the well was immersed in 0.4x SSC at 72°C for 2 minutes and 2xSSC + 0.05% Tween-20 at RT for 30 seconds. 100 μL of PBS was added to the well. The samples were imaged after 15 minutes.

#### Nascent-RNA labelling

Nascent RNA was labelled with Click-iT RNA Alexa Fluor 594 imaging kit (Thermo Fisher, C10330). SH-EP cells were plated on 8-well chambered coverglass (Thermo scientific 155411) and were infected with N-myc-mEGFP or N-myc/SparkDrop lentiviruses. Infected SH-EP cells were incubated with 1 µM lenalidomide for 2 hours to induce condensate formation. After labeled with 1 mM 5-ethynyl uridine (EU) for 1 hour, the cells were fixed with 4% paraformaldehyde in PBS, permeabilized with PBST (0.5% Triton X-100 in PBS), incubated with Click-iT reaction cocktail at room temperature for 30 mins to visualize 5-ethynyl uridine.

#### Western blot and estimation of protein levels

For detecting N-myc protein levels, cells were grown on 24 well plates, washed once with PBS, lysed with 100 μL lysis buffer (Cell signaling 9803S) at room temperature for 1 minute, mixed with 37.5 μL NuPAGE LDS sample buffer (NP0007) and 15 μL NuPAGE sample reducing agent (NP0009) and incubated at 70°C for 10 minutes. Protein samples were resolved on NuPAGE 4-12% Bis-Tris gel (NP0336), then transferred to nitrocellulose membrane under 25 V for 45 min in a Bio-rad Trans-Blot Turbo machine. The membrane was then blocked in 5% non-fat milk at room temperature for 1 hour, incubated with a MYCN mouse monoclonal antibody (Santa Cruz sc-53993, 2000X) at 4°C overnight, washed 3 times in PBST (0.1% Tween-20 in PBS), each time for 5 minutes, incubated with an HRP-conjugated anti-mouse secondary antibody (Cell signaling 7076S, 5000X) and washed 3 times, each time for 5 minutes. HRP chemiluminescent substrate (Thermo scientific 34580) was added to the membrane, incubated at room temperature for 5 minutes, then imaged using Bio-rad Chemidoc XRS system or film (Prometheus 30-507L) exposure. For β-actin, the staining process was similar, but the primary antibody (Santa Cruz sc-47778, 5000X) was incubated for 1 hour at room temperature.

To estimate the endogenous N-myc protein concentration in Kelly cells, we compared Kelly cells with the stable SH-EP cells expressing N-myc-mEGFP (SH-EP-MYCN) (See Supporting Fig. S5). First, we conducted western blot to compare the protein levels. ∼1000 cells were used for western blot. The bands were analyzed using ImageJ. Second, Kelly and SH-EP-MYCN were grown in glass-bottom 8-well chambers (Thermo Scientific 155411), stained with 1 μg/mL Hoechst 33342 for 10 minutes, and imaged under the confocal microscope for Hoechst fluorescence with the 100X objective. The nucleus was sectioned into multiple slices with a 0.5 μm interval. The nucleus volume was calculated using the 3D Objects Counter function in ImageJ. Third, SH-EP-MYCN cells were imaged in the GFP channel. The GFP fluorescence intensity was used to calculate N-myc-mEGFP concentration in the nucleus of the SH-EP-MYCN cells, by comparing its fluorescence intensity with that of purified mEGFP fluorescence intensity and concentration (Supporting Fig. S4). The Kelly cell N-myc concentration per nucleus was estimated by comparing the western-blot band intensity (Pillai-Kastoori et al., 2020), nuclear volume of Kelly vs SH-EP-MYCN cells, N-myc-mEGFP concentration in the nucleus of SH-EP-MYCN cells.

#### RT-qPCR

2×10^5^ of N-myc/SparkDrop and N-myc/SparkDrop control SH-EP stable cells were seeded in 24-well plate. 24 hours after seeding, the cells were challenged with 1 µM lenalidomide or 0.1% DMSO for 16 hours. The total RNA was extracted using a Quick-RNA Microprep Kit (Zymo Reserch, R1051) and converted to complimentary DNA using a SuperScript™ IV Reverse Transcriptase (Invitrogen, 18090010). The RT-qPCR was carried on a CFX96 Touch Real-Time PCR Detection System using iTaq Universal SYBR Green Supermix (Bio-Rad, 1725121) with a GAPDH control. The following primers were used.

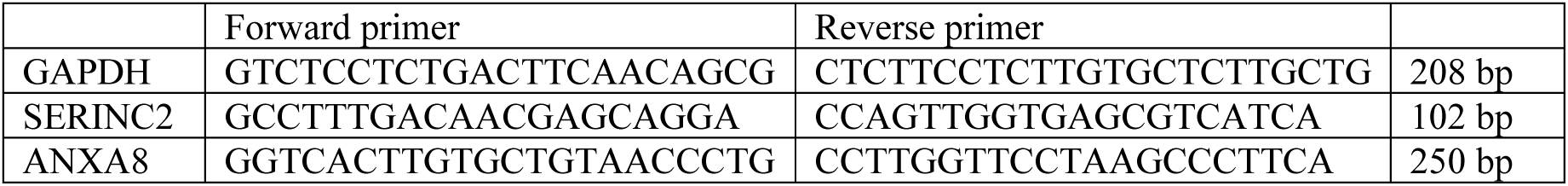

#### RNA sequencing and transcriptome analysis

SH-EP cells were infected with lentivirus expressing mEGFP as a baseline sample (without N-myc expression). N-myc/SparkDrop and N-myc/SparkDrop control (no HOTag6) SH-EP stable cells were seeded 1×10^5^ cells per cm^2^. 24 hours after seeding, the cells were treated with 1 µM lenalidomide or 0.1% DMSO for 16 hours. The total RNA was extracted using a Quick-RNA Microprep Kit (Zymo Reserch, R1051). The mRNA library construction and Illumina Novaseq sequencing were performed (Novogene Bioinformatics Technology Co., Ltd). 20 millions of 150 bp paired-end reads per sample were generated. After standard illumina Hiseq demultiplexing, raw data were processed with fastp (Chen et al., 2018) to remove PCR duplications and cut illumina adaptors. STAR (Dobin et al., 2013) was used to map reads to the human genome (hg38) by default setting. Then the raw counts of each gene were obtained. To identify differentially expressed genes (DEGs), edgeR (Robinson et al., 2010) was applied to the raw counts. Log2 Fold Change (log_2_FC), P value and false discovery rate (FDR) were calculated using TMM normalization. Then volcano plots were introduced to visualize DEGs analysis results (P < 0.01, FDR < 0.1) upon N-myc LLPS. The DEGs of N-myc in dilute phase (or condensed phase) to GFP baseline were determined by P < 0.01, FDR < 0.05 and |log_2_FC| > 0.3. Two groups of DEGs (N-myc dilute phase vs N-myc LLPS) were compared and those genes presented in both groups were considered as overlapped genes. Then Venn diagrams were generated by custom R scripts. Gene heatmap was plotted using differentially expressed genes by TPM normalization. Gene Ontology (GO) analysis of the LLPS-regulated DEGs were performed with ClusterProfiler (Robinson et al., 2010) for each library. E-box motif (CANNTG) in the DEGs promoter regions (3kb upstream) was detected by FIMO (v5.4.1) (Grant et al., 2011), then custom scripts was applied to calculate the count of E-box per gene. Wilcoxon test p-values were calculated for the E-box count between the LLPS-independent genes and LLPS-regulated genes (strong, weak).

---

**Fig. S1.**
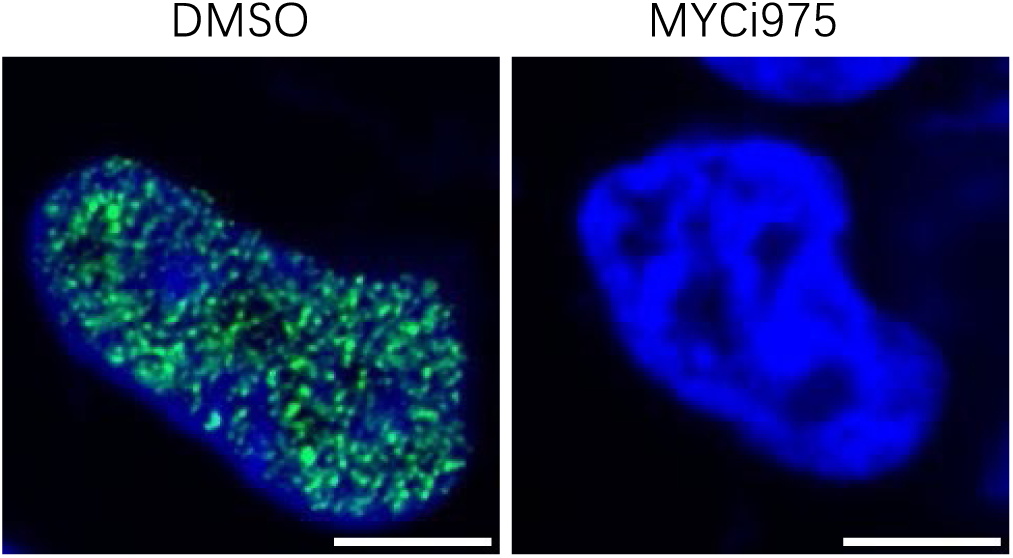
Immunofluorescence images of Kelly cells treated with DMSO or MYC/MAX dimerization inhibitor MYCi975. Kelly cells were incubated with DMSO or MYCi975 (30 μM) for 20 hours, followed by immunostaining with N-myc antibody. Scale bar, 5 μm.

**Fig. S2.**
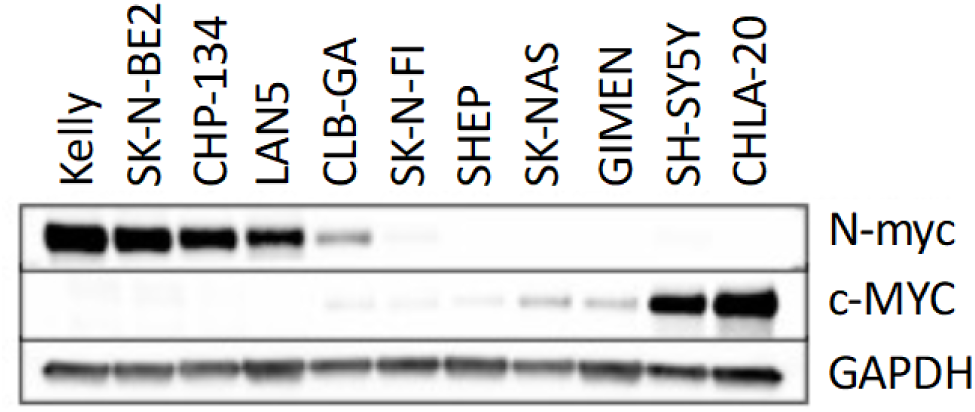
Neuroblastoma cell lines with various degrees of MYC (N-myc, c-MYC) protein expression. Western blot analysis of MYC proteins in various neuroblastoma cells.

**Fig. S3.**
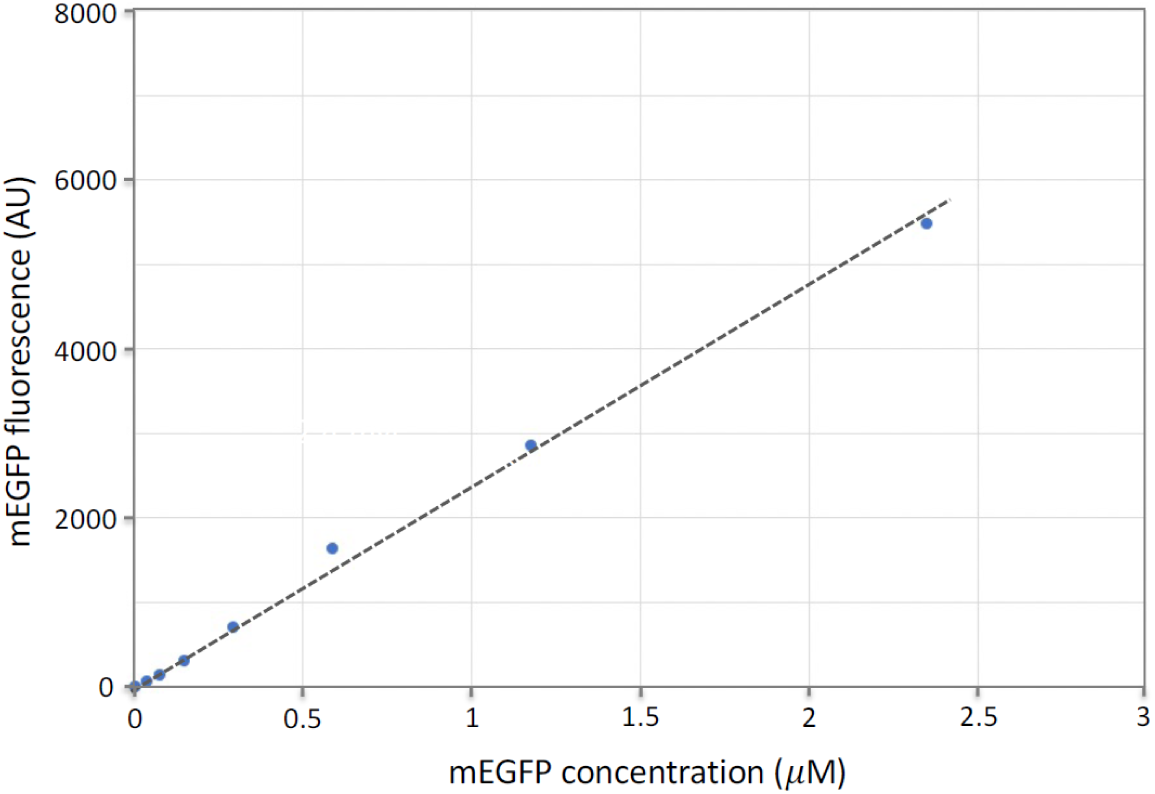
Relationship of mEGFP fluorescence brightness and concentration. mEGFP was purified and aliquoted at various concentrations. The protein samples were then imaged under the confocal microscope. The fluorescence brightness was recorded, corresponding to mEGFP concentration. The plotted line was used to estimate mEGFP concentration in living cells based on the fluorescence brightness.

**Fig. S4.**
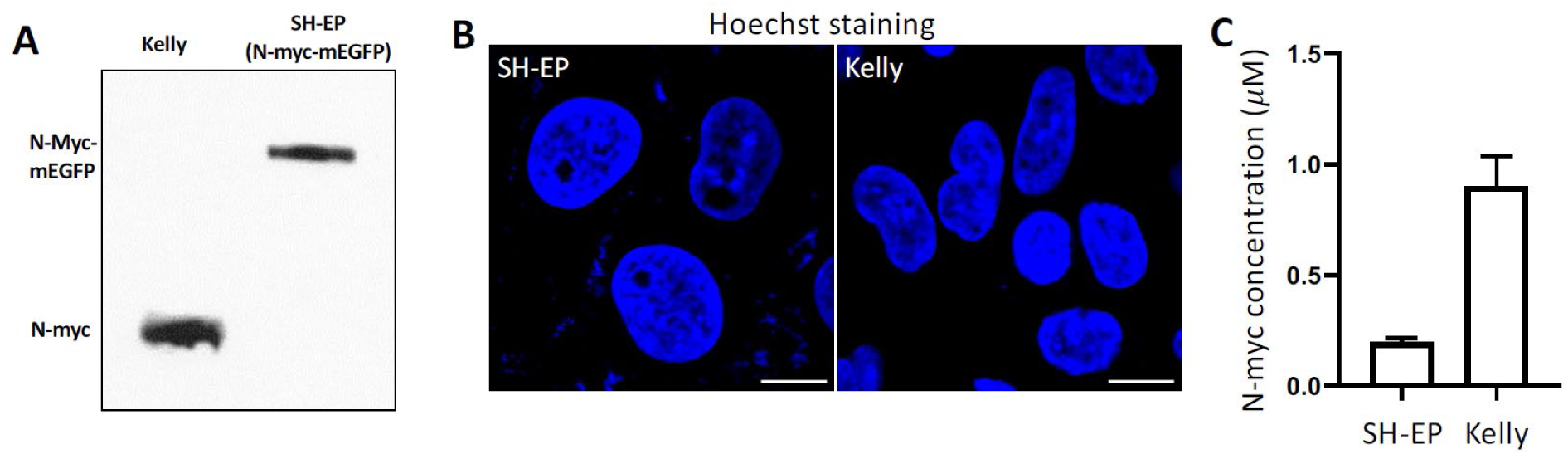
Estimation of N-myc concentration in the Kelly cells. The overall procedure is A) western blot analysis to determine relative amount of N-myc in Kelly vs SH-EP cells. B) Fluorescence imaging (Hoechst)-based measurement of the nuclear volume of SH-EP and Kelly cells. The N-myc-mEGFP concentration in SH-EP cells is determined by comparing the fluorescence intensity of N-myc-mEGFP versus that of the purified mEGFP. C) Estimation of the N-myc concentration in the nucleus of Kelly cells. Data are mean + SD (n = 20 cells). Scale bar, 10 μm.

**Fig. S5.**
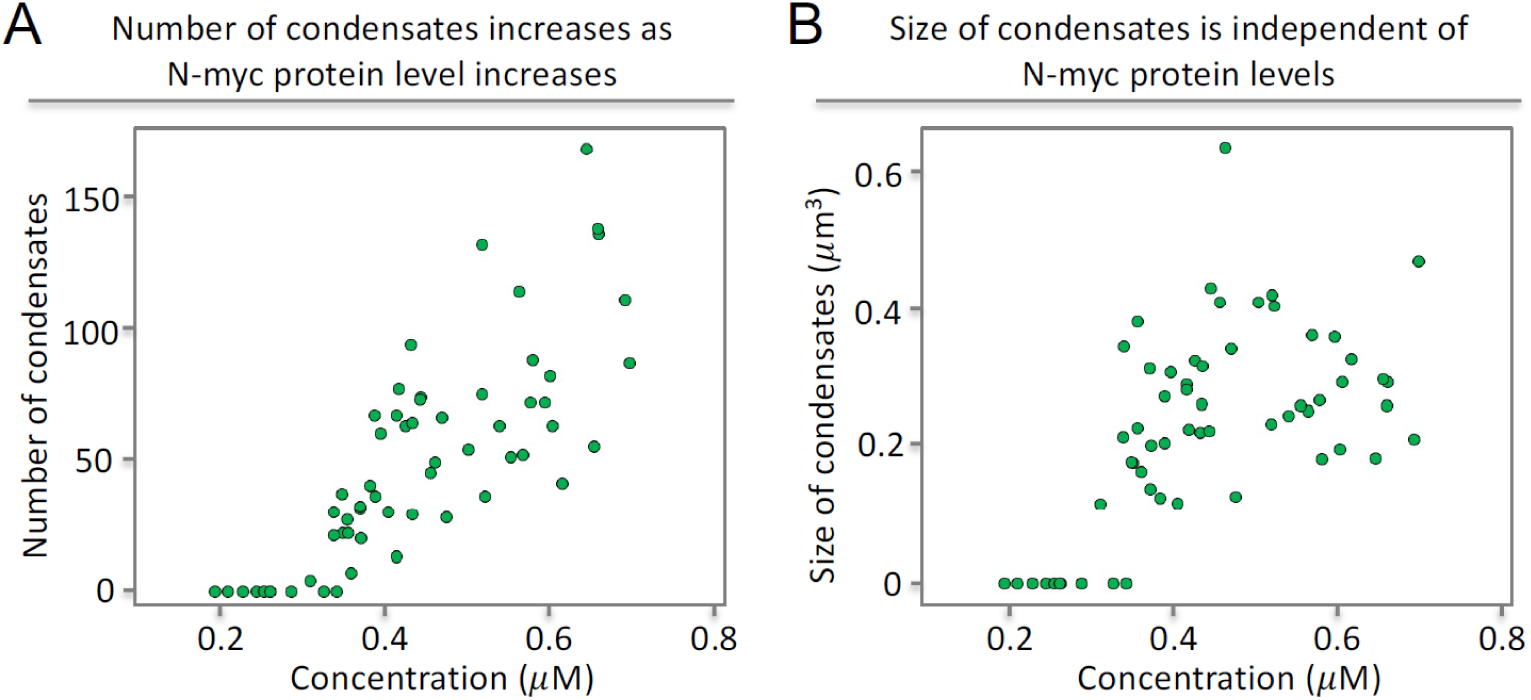
Relationship of N-myc condensate number and size to the N-myc protein levels. (A) Number of N-myc condensates was plotted against the N-myc protein level per nucleus. SH-EP cells expressing N-myc-mEGFP were imaged under the spinning disc confocal microscope. (B) Size of N-myc condensates was plotted against the N-myc protein level in single nucleus.

**Fig. S6.**
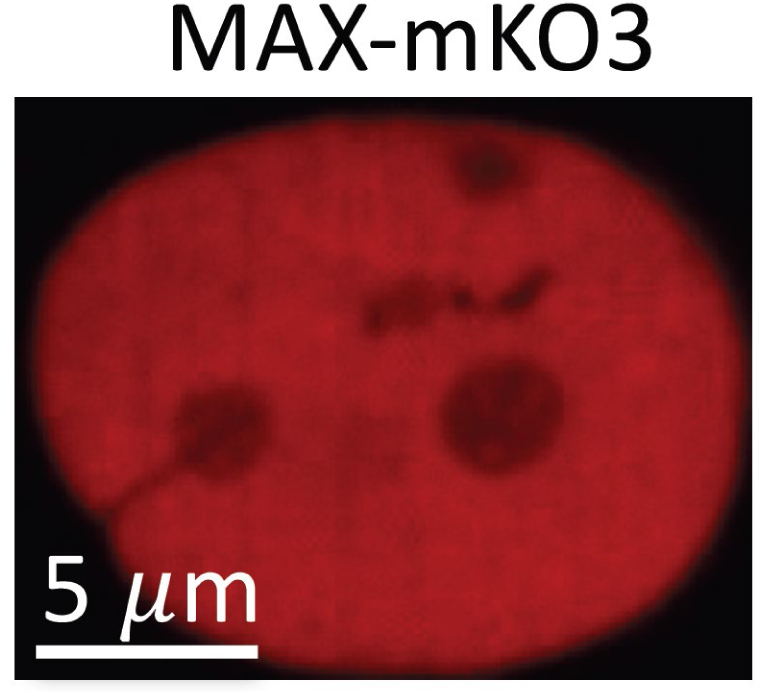
MAX does not form condensates in the absence of N-myc condensates. Fluorescence imaging of the nucleus of a SH-EP cell expressing MAX-mKO3. Scale bar, 5 µm.

**Fig. S7.**
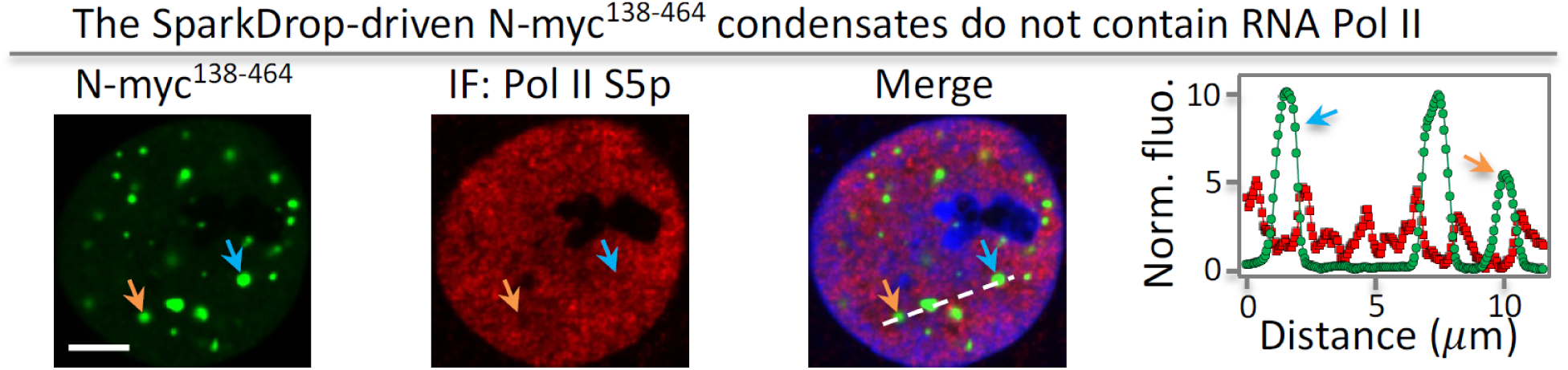
The induced condensates of TAD-truncated N-myc mutants do not contain RNA polymerase II. Immunofluorescence images of Pol II S5p with N-myc^138–464^/SparkDrop condensates. Scale bar, 5 μm.

**Fig. S8.**
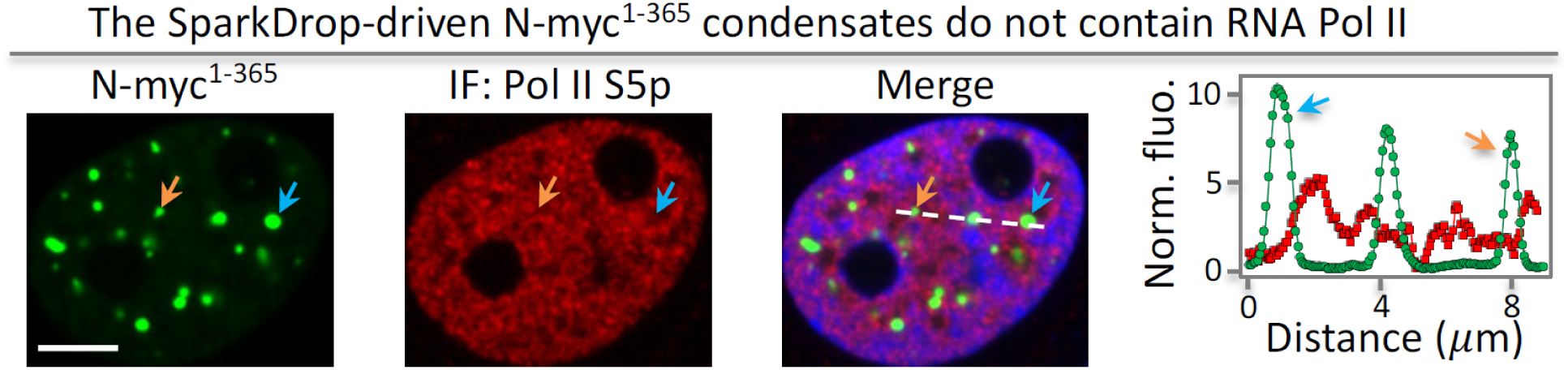
The induced condensates of C-terminal truncated N-myc mutants do not contain RNA polymerase II. Immunofluorescence images of Pol II S5p with N-myc^1–365^/SparkDrop condensates. Scale bar, 5 μm.

**Fig. S9.**
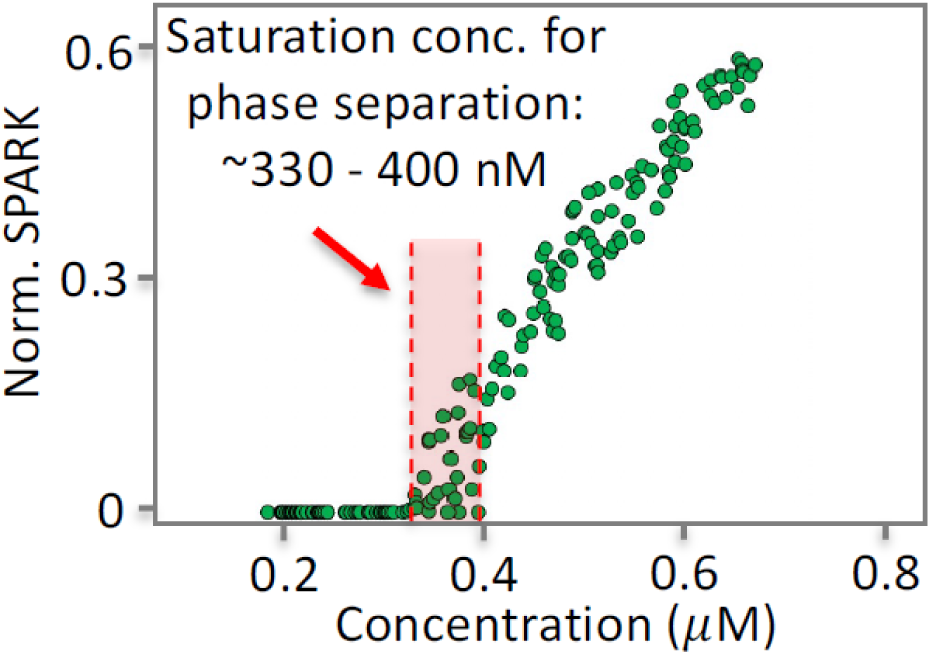
SparkDrop-tagged N-myc phase separates with threshold concentration around 330 to 400 nM. SparkDrop-tagged N-myc was expressed in SH-EP cells. Each green dot represents data from a single cell (∼200 cells). The concentration was calculated based on the green fluorescence of mEGFP in SparkDrop.

**Fig. S10.**
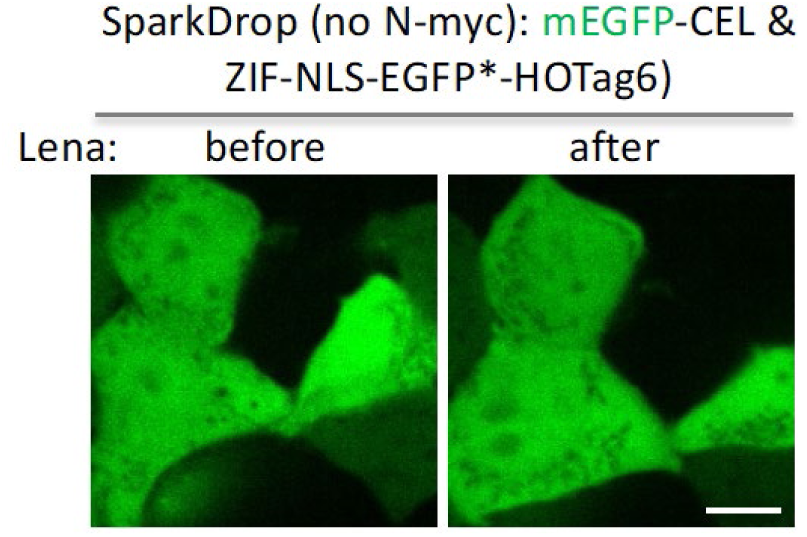
SparkDrop without N-myc does not form condensates upon addition of lenalidomide. SH-EP cells expressing the SparkDrop without N-myc were treated with lenalidomide. Scale bar, 10 μm.

**Fig. S11.**
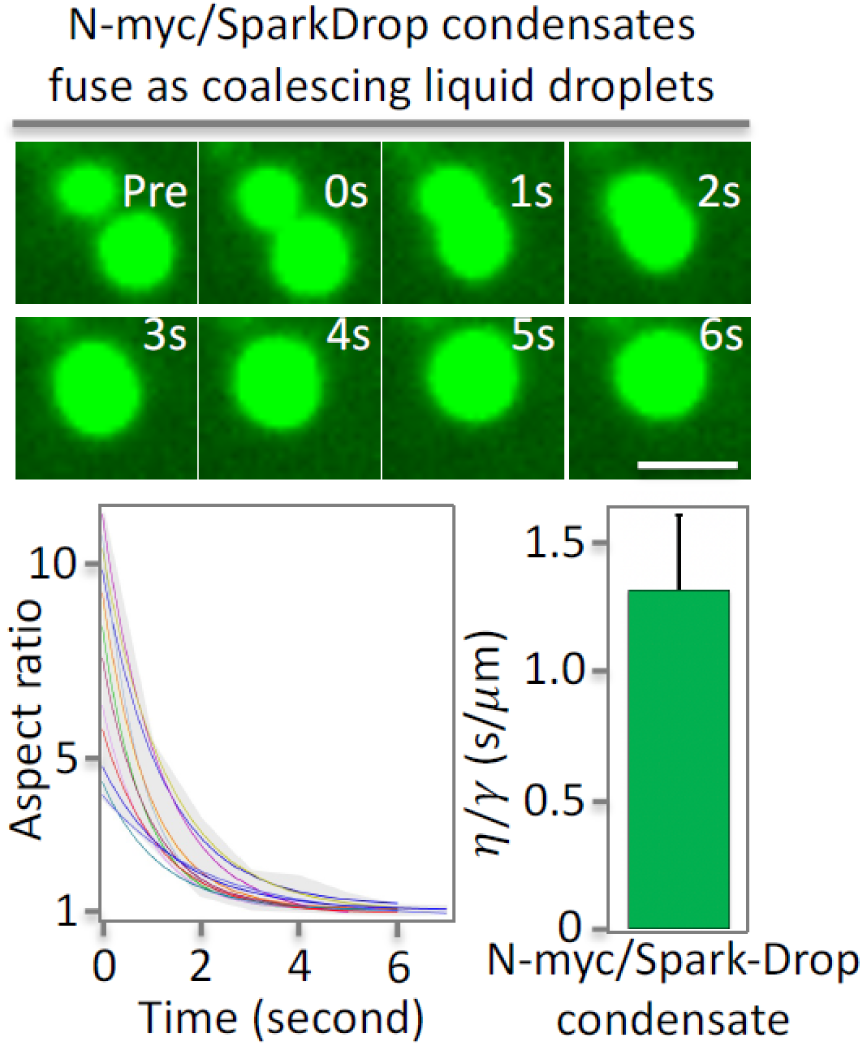
SparkDrop-tagged N-myc condensates are liquid droplets. Fusion events between N-myc/SparkDrop condensates. Top: fluorescence images. Bottom-left: quantitative analysis of the fusion events shown in the top. Bottom-right: inverse capillary velocity. Error bar represents standard deviation (n = 12). Inverse capillary velocity is 1.31 ± 0.29 (s/μm). Scale bar, 1 μm.

**Fig. S12.**
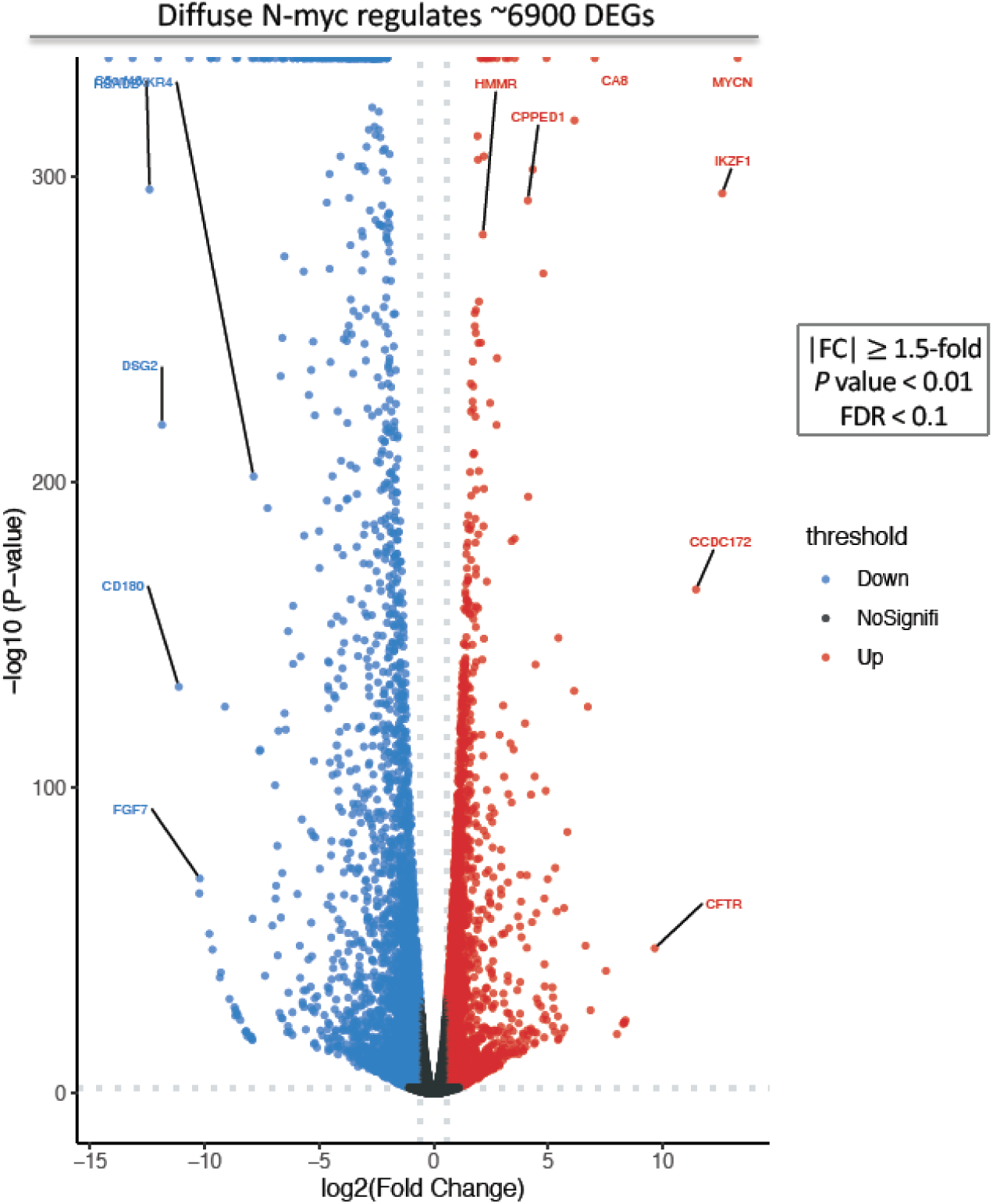
Volcano plot showing DEGs that are regulated by diffuse N-myc (no phase separation). SH-EP cells expressing N-myc/SparkDrop in diffuse state were processed for RNA sequencing. mRNAs showing significant up- and down-regulation (|FC| ≥ 1.5, p-value < 0.01, FDR < 0.1) are marked in red and blue, respectively. Black dots represent mRNAs with no significant changes.

**Fig. S13.**
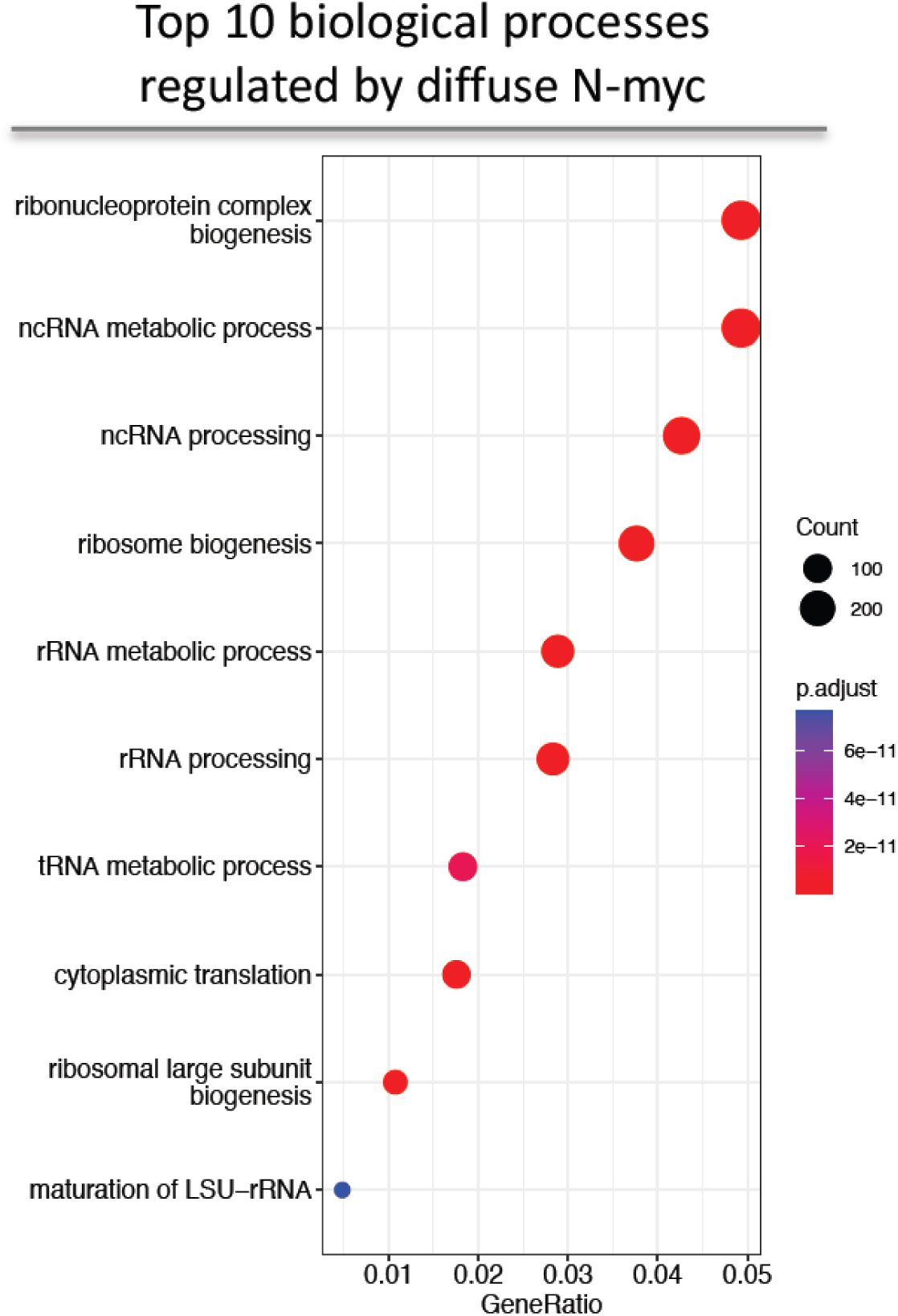
GO enrichment analysis of the biological processes that are regulated by diffuse N-myc (no phase separation). N-myc/SparkDrop-expressing cells without lenalidomide treatment were used for RNA sequencing and analysis.

**Table S1.**
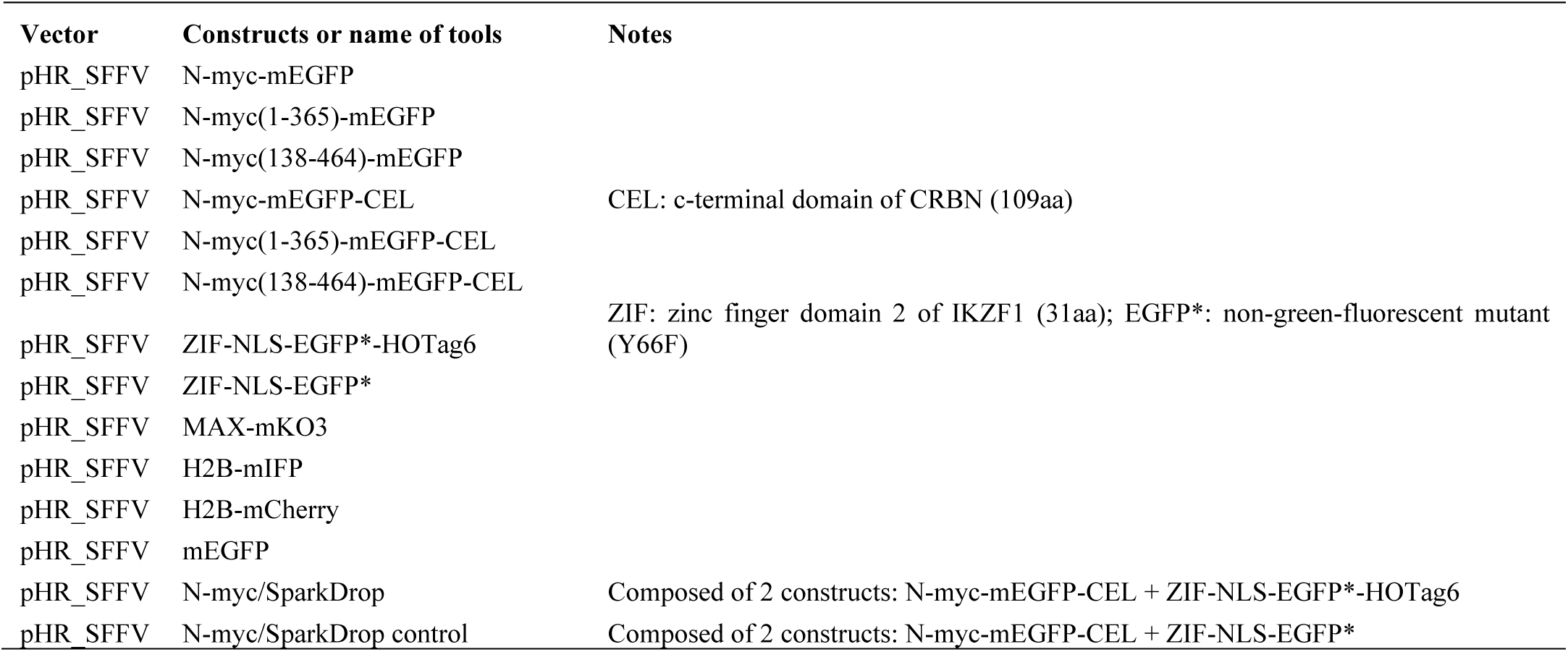
List of all constructs.

